# Molecular Origins of pH Gradients in Charge-Regulated Biomolecular Condensates

**DOI:** 10.64898/2026.07.02.736097

**Authors:** Shuo-Lin Weng, Shiv Rekhi, Young C. Kim, Jeremy C. Palmer, Jeetain Mittal

## Abstract

Biomolecular condensates exhibit spontaneous electrochemical microenvironments characterized by asymmetric ion distributions and pH gradients that emerge from protein-sequence-dependent charge regulation. Despite their biological importance, mechanistic understanding of these microenvironments has been constrained by the absence of computationally tractable frameworks capable of treating proton exchange, counterion partitioning, and buffer equilibria on consistent thermodynamic footing. Here, we introduce the buffered Charge-Regulation Monte Carlo (b-CR-MC) framework, which couples grand-canonical exchange of ions and buffer species with explicit charge regulation of titratable residues. By extending the CR-MC ion-merging strategy to multicomponent reservoirs and employing the Restricted Primitive Model, b-CR-MC achieves computational efficiency while maintaining thermodynamic rigor, with quantitative agreement to the more expensive generalized G-RxMC approach. Applied to full-length FUS (net positive) and PGL-3 (net negative) under physiological conditions, the framework reveals sequence-dependent pH gradients: the dense phase of FUS exhibits an alkaline shift, while PGL-3 exhibits an acidic shift, in both cases driving the condensate interior toward the protein’s isoelectric point. Slab-geometry simulations further resolve the Donnan potential and continuous ion profiles across the condensate interface, confirming the direction and magnitude of these electrochemical shifts. Additionally, we identify spatially resolved buffer depletion within dense phases, establishing that dynamic charge regulation is a primary determinant rather than a secondary correction to condensate electrochemistry. By establishing a sequence-resolved, thermodynamically consistent computational platform, b-CR-MC enables quantitative prediction of how mutations and post-translational modifications reprogram condensate microenvironments across biological and pathophysiological contexts.

## I. INTRODUCTION

Biomolecular condensates, membraneless organelles formed by the liquid–liquid phase separation (LLPS) of intrinsically disordered proteins and nucleic acids, are increasingly recognized as active biochemical regulators rather than passive storage depots.^1^ Their dense interiors host non-uniform ion distributions, interphase electric potentials, and spontaneous pH gradients that differ substantially from the surrounding cellular environments.^2,3^ These specialized electrochemical microenvironments reshape macromolecular interaction networks, modulate sequestered enzyme activity, and influence post-translational modifications.^4^ Consequently, pH- and ion-dependent changes in macromolecular charge state feed back onto phase behavior, altering condensate material properties, compositions, and cellular persistence, a coupling with direct implications for disease-associated condensates such as those formed by the ALS-linked RNA-binding protein FUS.^5^ A molecular-level understanding of how such electrochemical features emerge from protein sequence is therefore central to explaining condensate physiology and pathology.

Extensive theoretical and computational work with coarse-grained (CG) models has established that sequence-encoded properties, including net charge, charge patterning, and hydropathy, strongly influence condensate stability and composition.^6–10^ However, most existing studies employ a fixed-charge approximation, assigning each ionizable residue a static charge state based on its dilute-solution pKa and an externally imposed pH.^11–13^ This approximation implicitly assumes that the dense-phase dielectric environment, macromolecular crowding, and the Donnan potential arising from asymmetric ion partitioning do not significantly perturb acid–base equilibria inside condensates. In reality, altered solvation, fluctuating counterion concentrations, and interchain electrostatics shift pKa values relative to bulk solution, leading to changes in both net charge and charge patterning upon condensation that fixed-charge models cannot capture self-consistently.^14–17^ Experimental measurements demonstrate that condensate dense phases can sustain pH values that spontaneously shift toward the protein isoelectric point, a charge neutralization effect tunable via charged client recruitment.^3^ All-atom constant-pH (cpH) molecular dynamics (MD) simulations further support dense-phase-induced pKa shifts.^17^ Together, these observations establish a fundamental coupling among charge regulation, ion partitioning, and condensate-internal pH that demands a grand-canonical theoretical framework to be resolved at the sequence level.

The appropriate theoretical setting for this problem is the grand-canonical ensemble, in which titratable residue protonation states, counterion numbers, and buffer composition all fluctuate in equilibrium with a thermodynamically defined reservoir.^18,19^ Standard cpH methods struggle with this requirement; enforcing strict electroneutrality demands precise synchronization between titration and explicit ion exchange events, while uncorrected violations distort electrostatic potentials and Donnan equilibria.^19–21^ Standard cpH implementations are further limited by systematic dense-phase pH errors, high computational costs, or convergence difficulties at condensate scales.^20–24^ Grand-Reaction Monte Carlo (G-RxMC), introduced by Landsgesell et al.,^18^ couples reactive acid–base sampling to grand-canonical ion exchange via reaction ensemble algorithm,^25–27^ capturing Donnan partitioning and electrostatic correlations, and has been applied to ionization equilibria in polyelectrolyte brushes, gels, and complexes.^16,28–34^ Its generalized variant, g-G-RxMC, developed by Beyer and Holm, extends this capability to complex reservoirs by allowing weak polyprotic acid buffer species to exchange.^35^ In parallel, the Charge-Regulation Monte Carlo (CR-MC) method of Curk, Yuan, and Luijten streamlines this reaction framework by merging monovalent ions under the Restricted Primitive Model (RPM), in which ions of the same valency share identical size and interaction parameters.^36^ Despite this progress, no existing framework has been applied to sequence-resolved protein condensates, where heterogeneous pKa values, complex charge patterning, and dense macromolecular environments jointly shape electrochemical behavior, leaving the molecular origins of condensate pH gradients and sequence dependence of buffer partitioning unresolved.

To close this gap, we introduce the buffered Charge-Regulation Monte Carlo (b-CR-MC) framework and apply it to sequence-resolved CG simulations of condensates formed by full-length FUS and PGL-3. By extending the CR-MC ion-merging strategy to multicomponent buffered reservoirs and demonstrating its thermodynamic equivalence to g-G-RxMC under the RPM, b-CR-MC enables efficient grand-canonical sampling of charge regulation, ion partitioning, and buffer exchange in sequence-resolved protein systems. We validate the framework against g-G-RxMC in protein-free box and single-chain limit. Using dual-box phase coexistence simulations, we demonstrate that FUS and PGL-3 condensates support oppositely directed pH gradients, each directed toward the respective protein isoelectric point, accompanied by distinct patterns of buffer depletion and charge redistribution invisible to fixed-charge models. A slab-geometry simulation of FUS independently resolves the continuous Donnan potential and ion profiles across an explicit condensate interface, corroborating the alkaline pH shift obtained from the dual-box protocol. Together, these results establish charge regulation and buffer partitioning as primary determinants of condensate electrochemical microenvironments, demonstrating that b-CR-MC provides a practical, thermodynamically rigorous platform for connecting protein sequence to condensate electrochemical function.

## II. THEORETICAL AND COMPUTATIONAL METHODS

### A. Buffered Charge-Regulation Monte Carlo (b-CR-MC)

Charge regulation in biomolecular condensates requires a grand-canonical treatment in which protonation states, ion numbers, and buffer composition all fluctuate in equilibrium with a macroscopic reservoir. The b-CR-MC framework achieves this by extending the ion-merging strategy from the original CR-MC approach of Curk et al. to multicomponent buffered reservoirs^36^. Under the Restricted Primitive Model (RPM), in which same-valency ions share identical mass, size, and interaction parameters, all monovalent cations are merged into a single species X^+^ = {H^+^, Na^+^} and all monovalent anions into X^−^ = {OH^−^, Cl^−^}. This ion-merging is thermodynamically exact: the grand-canonical partition function of b-CR-MC is identical to that of the generalized Grand-Reaction Monte Carlo (g-G-RxMC) formulation of Beyer and Holm under the RPM^35^. The full reaction set and the derivation of thermodynamic equivalence are provided in **Supplementary Material, Section S1**. The mapping reduces the reaction pool to 3 + 2N entries for a system with N titratable residue types, a reduction of more than threefold that scales favorably with sequence heterogeneity. The mapping introduces two composite quantities, encoding the sodium-to-proton fugacity ratio:

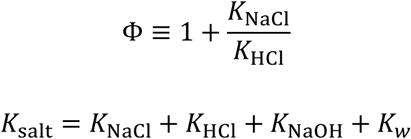

The complete b-CR-MC reaction set comprises three classes, where Δ*ν* denotes the net change in the number of particles in the forward reaction:

1. Ion and buffer exchange reactions:

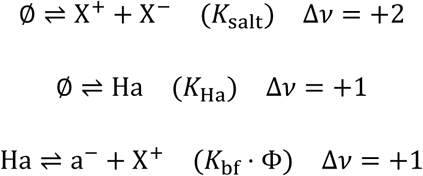

2. Titratable residue reactions:

Acidic (e.g., Asp, Glu, Tyr):

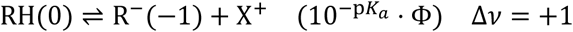

Basic (e.g., Lys, Arg, His):

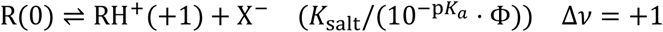

For basic residues, protonation of the neutral state R(0) is coupled to an ion-pair insertion move rather than an explicit deprotonation reaction, ensuring strict charge neutrality throughout the simulation.

3. Swap moves: Each residue type retains an identity-swap reaction (Kswap = 1) to accelerate protonation-state mixing within a given titratable type.

### B. Reaction ensemble algorithm

The reaction ensemble algorithm was employed to model both chemical reactions within the system and particle exchange with an external reservoir.^26,27^ The forward acceptance probability for a reaction *r* is:

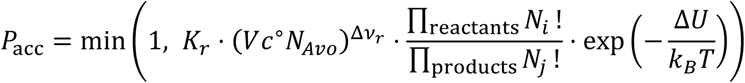

where Kr is the equilibrium constant of reaction *r* at standard concentration *c*^∘^ = 1 M, V is the simulation box volume, NAvo is Avogadro’s number, Δ*ν_r_* is the net change in the particle number for the trial move, Ni and Nj represent the particle numbers of the reactant species before the reaction move and the product species after the move, and Δ*U* is the change in potential energy.

### C. Monovalent buffer representation

In the simulations reported here, the buffer is modeled as an effective monovalent weak acid with pKa = 7.20, chosen to match the second dissociation constant of phosphoric acid (pKa2 = 7.20), which governs buffering capacity in the physiological pH range (∼7.0–7.5). This abstraction, treating the buffer as a generic acid-base pair rather than as an explicit phosphate species, is required to preserve compatibility with the RPM: inclusion of the divalent form (e.g., HPO_4_ ^2-^) would violate the RPM species-indistinguishability condition and thereby invalidate the thermodynamic equivalence derived above.

### D. Effective pH

The local pH in each phase was determined self-consistently from the Henderson–Hasselbalch relation applied to the sampled [Ha]/[a^−^] ratio:

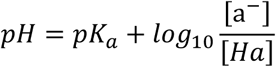

This expression assumes unit activity coefficients; in the non-ideal condensate environment, the thermodynamically exact pH would require excess chemical potentials for Ha and a⁻. Because these quantities are not directly accessible from a CG simulation without additional model-dependent assumptions, the Henderson–Hasselbalch estimate is employed as a practical, internally consistent approximation.^18,34,35,37^ All reported pH values should be interpreted as effective pH values that correctly reflect the local protonation equilibrium within the model.

### E. Assessment of the RPM approximation error

While the ion-merging approximation introduced by the RPM is exact when all same-valency ions share identical size and interaction parameters, its application harbors two principal limitations. First, in practice, monovalent cations differ in hydrated radius: H⁺ (∼2.8 Å) is smaller than Na⁺ (∼3.6 Å),^38^ introducing a size asymmetry that the merged X⁺ representation neglects; an analogous asymmetry exists for the merged X-species. An order-of-magnitude estimate based on Debye–Hückel theory indicates that this asymmetry perturbs the dilute-phase electrostatic screening length by a few percent relative to the symmetric-RPM value,^39^ leading to a slight deviation in dilute-phase ionization predictions. It remains highly conceivable, however, that dense-phase macromolecular crowding could amplify short-range ion–protein correlations beyond these ideal dilute-limit expectations. A systematic quantification of the RPM approximation error in protein-dense environments is therefore a priority for future benchmarking. Second, the monovalent buffer representation introduces an additional quantifiable limitation: within Donnan theory, the partition coefficient of an ion of valency z scales as *ξ* ∝ *exp*(−*zβψ_Don_*), so divalent anions are exponentially more sensitive to the Donnan potential than monovalent ones. At pH 7.4, where HPO_4_ ^2-^/H PO ^-^ ratio is approximately 4:1,^40^ the replacement of this predominantly divalent buffer with a monovalent surrogate is expected to affect Donnan-driven buffer exclusion. Furthermore, such a substitution risks disregarding critical multivalent interactions and intra-condensate accumulation, consistent with recent PEC observations.^32^ Consequently, all results reported here therefore pertain to a generic monovalent weak acid at pKa = 7.20; extension to multivalent buffer representations requires the full g-G-RxMC framework.^35^

### F. Chemical potential tuning (μ-tuning)

Because buffer species interact non-ideally within the simulation box, the equilibrium constants {Ki} encoding the target reservoir composition (150 mM NaCl, 12 mM buffer, pH 7.40) cannot be determined from the Henderson–Hasselbalch equation alone and must be calibrated iteratively. We employed the μ-tuning algorithm of Beyer and Holm,^35,41^ in which the effective chemical potential of species *j* at iteration *t* is updated as:

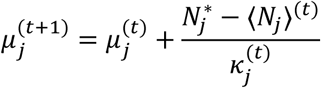

where 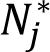 is the target number of particles, ⟨*Njj*⟩(t) is the ensemble average at iteration *t*, and 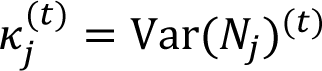 = Var(*N*) (t) is estimated from the second half of each iteration’s trajectory, serving as a variance-based step-size regulator. To ensure numerical stability during early iterations, 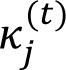 is bounded between *κ_min_* = 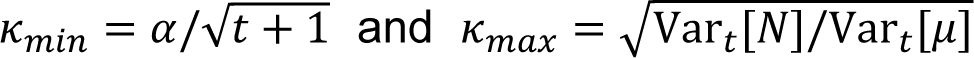 and *κ_max_* = 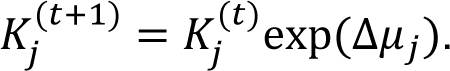 where *α* ∝ *V*/*U* with the system volume V and a characteristic energy scale U.^35^ The equilibrium constant is then updated as *K_j_*^(t+1)^ = *K_j_*^(t)^exp(Δ*μ_j_*). Convergence is declared when the relative deviation between the ensemble-averaged and target particle numbers falls below 1% for NaCl and 5% for buffer species. The resulting {K_i_} set was applied without further adjustment to all protein-containing simulations, ensuring that both the dense and dilute phase boxes are thermodynamically coupled to a common macroscopic reservoir.

### G. Hybrid MD/MC scheme in HOOMD-blue

Production simulations employed a hybrid Langevin dynamics–reactive Monte Carlo (MD/MC) scheme implemented within HOOMD-blue^42^ via a custom plugin developed as part of this work. The plugin transfers the functionality of the ConstantpHEnsemble and ReactionEnsemble modules from ESPResSo to HOOMD-blue, incorporating a virtual-particle strategy to realize grand-canonical insertions and deletions within HOOMD-blue’s natively canonical ensemble.^43,44^ Rather than creating or destroying particles on the fly, identity toggles switch particles between active (real) and non-interacting virtual states; virtual particles are uncharged and strictly non-interacting, making this protocol thermodynamically equivalent to explicit insertion/deletion while avoiding the associated bookkeeping overhead. Prior to each production run, a pool of virtual particles—sized to comfortably exceed the maximum anticipated real-particle count—was pre-allocated in the simulation box to ensure proper semi-grand canonical sampling. Plugin fidelity was verified against established reference results, with all benchmarks agreeing with ESPResSo to within statistical uncertainty.^34^ A demonstration of the thermodynamic equivalence of the virtual-particle strategy, details of the benchmarking systems, and the corresponding results are provided in **Supplementary Material**.

The equations of motion were integrated using a Langevin thermostat at 300 K with a time step of dt = 10 fs. Following every 1000 MD steps, 100 MC attempts were executed, with reactions selected uniformly at random from the pool and evaluated via the Metropolis criterion. This MD/MC ratio was applied uniformly across all systems.

### H. Coarse-grained model

Proteins were modeled at single-bead-per-residue resolution using the hydropathy-scale (HPS) Urry coarse-grained model,^6,9^ with altered electrostatic potential treatment described below. The parameters for titrated forms are scaled from the original hydropathy table of Urry et al.^45^ Backbone connectivity was enforced by harmonic bonds (k = 20 kcal/(mol Å^2^), r0 = 3.8 Å), while nonbonded residue–residue interactions were described by the Ashbaugh–Hatch (LJ-λ) potential with ε = 0.2 kcal/mol.^46^ Ions and buffer species (X^+^, X⁻, Ha, and a⁻) interacted with amino-acid beads and with each other through purely repulsive LJ-λ potentials, equivalent to the Weeks-Chandler-Andersen (WCA) potential (λ = 0, ε = 1.0 kcal/mol), with size parameters determined by Lorentz–Berthelot combining rules.^47^ All merged ions (X⁺ and X⁻) were assigned a common diameter σ = 3.55 Å, consistent with the widely used polymer models.^18,32,34–36^ This uniform assignment preserves the RPM indistinguishability condition for both ion–ion and ion–protein interactions. Electrostatic interactions between all charged residues, ions, and buffer species were computed using the particle–particle particle-mesh (PPPM) method as implemented in HOOMD-blue.^42,48^ The unit charge was scaled to 2.037 to include the prefactor of 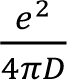 under a dielectric constant D of 80 for water.

Six amino-acid types were treated as titratable in the simulations: Asp (pKa 3.67), Glu (4.25), His (6.54), Tyr (9.84), Lys (10.53), and Arg (12.48). The simulated sequences are listed in **Table S1**, and parameters for titratable residues and ion species are listed in **Table S2**. Folded domains were constrained as rigid bodies using a custom plugin that enforces structural rigidity while permitting constituent residues to titrate freely: the RNA recognition motif (RRM, residues 285–371) and zinc finger (ZnF, residues 414–454) of FUS, and to the N-terminal (NtDD, residues 1–205) and central (CeDD, residues 206–447) dimerization domains of PGL-3. While this rigid-body treatment preserves structural integrity, it explicitly decouples charge regulation from conformational fluctuations within these domains. Future studies should address how this decoupling impacts local electrostatic landscapes and condensate phase behavior.

### I. Phase coexistence simulations

Two complementary protocols were used to characterize condensate electrochemistry.

#### Dual-box protocol

Dense and dilute phases were represented by separate simulation cells, following the approach of Staňo et al.,^32^ thermodynamically coupled to a common reservoir via identical μ-tuned equilibrium constants {Ki} to both simulation boxes. These constants simultaneously impose 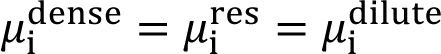 for all exchangeable species, ensuring that observed electrochemical differences between boxes reflect the intrinsic thermodynamic response of each phase rather than parametric drift.

The dense-phase box contained 50 protein chains in a 22 nm cubic cell, yielding a protein concentration of approximately 7.8 mM. The equilibrium box dimensions were determined by a preceding 100 ns NPT relaxation at zero applied pressure in the absence of reactive Monte Carlo updates, following the established protocol.^8^ The dilute-phase box contained a single protein chain in a 30 nm cubic cell, a volume sufficiently large to preclude artificial self-interactions across periodic boundaries. All production MD/MC simulations were subsequently performed at fixed temperature and volume.

Because protein chains were not exchanged between boxes, this protocol simulates the condensate interior at a prescribed density rather than capturing true thermodynamic phase coexistence. Nonetheless, the resulting electrochemical observables, including pH, ion partitioning, and charge regulation, accurately reflect the thermodynamic response of each phase at fixed density, a standard convention in biomolecular condensate simulations.^6,8^ A defining feature of this framework is that the interphase Donnan potential is not pre-parameterized; instead, it emerges self-consistently from the grand-canonical charge-regulation moves.^16,18,35,37^ Each reactive Monte Carlo step dynamically adjusts local protonation states and ionic composition to satisfy both reservoir chemical potentials and the instantaneous electrostatic environment, ensuring that the distinct ion and buffer distributions between the dense and dilute boxes represent a self-consistent thermodynamic response to different macromolecular environments.

#### Slab-geometry protocol

To resolve electrochemical profiles across a continuous condensate interface, we simulated the direct phase coexistence of FL-FUS using a slab geometry adapted from established protocols.^6,9^ The initial slab configuration was prepared by placing 40 chains of the FL-FUS monomeric sequence within an elongated simulation cell (15 × 15 × 105 nm). Utilizing the identical μ-tuned equilibrium constants {Ki}, the slab system was thermodynamically coupled to the same macroscopic reservoir (150 mM NaCl, 12 mM buffer, pH 7.40) used in the bulk simulations. Unlike the dual-box method, the slab geometry permits the spontaneous formation of a dense liquid phase in direct physical contact with the surrounding dilute phase within a single simulation volume. This explicit interface enables the grand-canonical b-CR-MC framework to continuously map spatial variations in ion partitioning, buffer distribution, and sequence-specific charge regulation directly across the dense–dilute phase boundary.

## III. RESULTS AND DISCUSSION

### A. A Transferable Thermodynamic Reference for Grand-Canonical Condensate Simulations

Meaningful two-phase electrochemical measurements require that the chemical potentials of all exchangeable species be precisely encoded before any protein is introduced. In the dual-box protocol, both the dense and dilute phases are thermodynamically coupled to a common macroscopic reservoir via a shared set of equilibrium constants {Ki}; any inaccuracy in these constants propagates directly into the partitioning of ion and buffer species, biasing the computed Donnan potential, the interphase pH gradient, and all charge-regulation observables. We first tested whether the iterative μ-tuning procedure could faithfully recover the target reservoir composition (150 mM NaCl, 12 mM monovalent buffer, pH 7.40) without protein-specific parameter adjustment.

The equilibrium constants were calibrated by the μ-tuning procedure using the standard g-G-RxMC reaction pool (10 reactions, corresponding to N = 0 titratable residue types, see **Methods**) in a protein-free 20 nm cubic box, corresponding to target particle counts of approximately 758 Na^+^, 723 Cl^-^, 22 Ha, 35 a^-^, 0 H^+^, and 0 OH^-^. Starting from an initial analytical estimate, the iterative algorithm converged within eight cycles: the ensemble-averaged particle counts for all ionic and buffer species stabilized within the prescribed tolerances, and the self-consistent pH, computed from the Henderson–Hasselbalch relation applied to the converged 〈*N*Ha〉/〈*N*a− 〉 ratio, reproduced the target value of 7.40 to within numerical precision (**Fig. 1**).

**Fig. 1.**
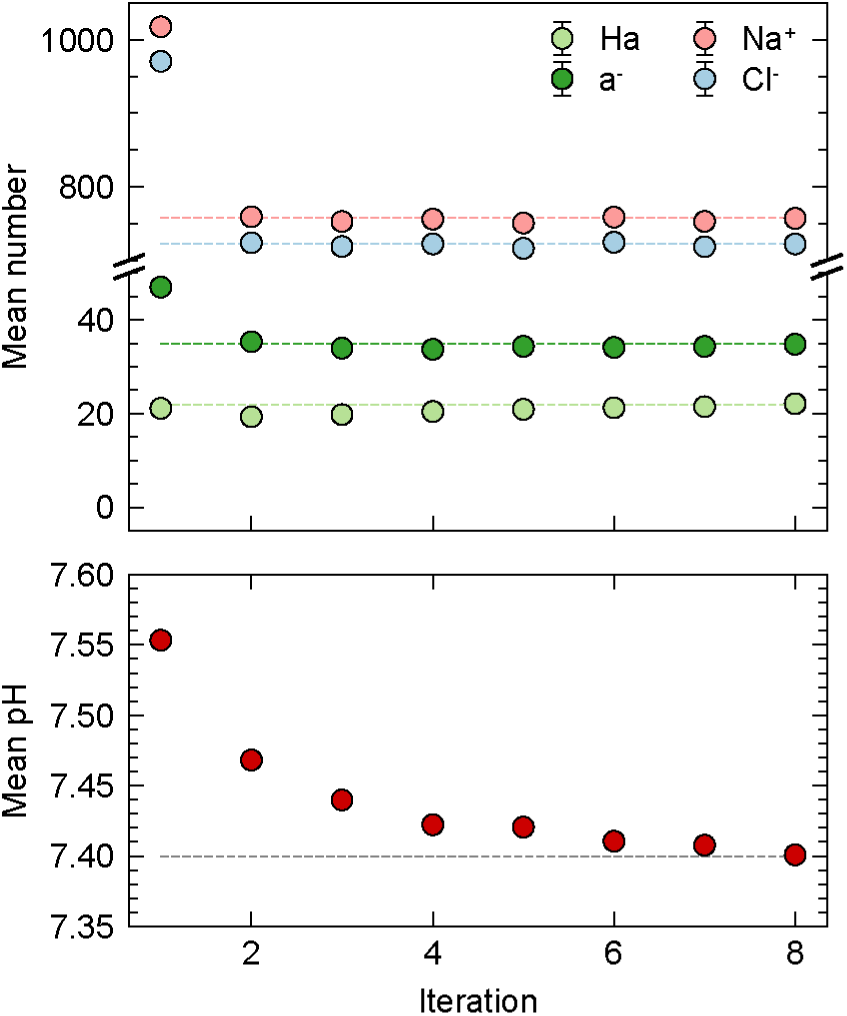
Convergence of reservoir properties during chemical potential μ-tuning. (Top) Ensemble-averaged particle counts for buffer species (Ha, a⁻) and salt ions (Na⁺, Cl⁻) as a function of tuning iteration. Dashed lines indicate the target particle numbers for a 20-nm cubic simulation box. **(Bottom)** Corresponding self-consistent mean pH across iterations. The system rapidly converges to the target macroscopic conditions (12 mM buffer, 150 mM NaCl, pH 7.40) within eight iterations.

The resulting {Ki} set serves as a transferable thermodynamic reference that is applied without further adjustment to all subsequent protein-containing simulations. This invariance ensures that any electrochemical differences observed between the dense and dilute phases reflect the intrinsic response of the macromolecular environment rather than parametric drift. This transferability thereby establishes the rigorous baseline required to compare electrochemical behavior across proteins with disparate sequence compositions.

### B. b-CR-MC Preserves Thermodynamic Accuracy Relative to the Full Multi-Species Formulation

The foundational approximation of b-CR-MC is that merging monovalent ions into composite species X^+^ and X^-^ under the RPM reduces the reaction pool without altering thermodynamic observables. To confirm that this reduction introduces no measurable bias in either bulk buffer equilibria or sequence-specific protein ionization, we benchmarked b-CR-MC against the explicit g-G-RxMC formulation at two levels of complexity: a protein-free reservoir box and a single-chain simulation of full-length FUS (FL-FUS) at pH 7.40.

In the protein-free box, both methods yield indistinguishable ensemble-averaged populations for Ha and a^-^, with self-consistent pH values agreeing to within numerical precision, demonstrating that the merged X^±^ representation reproduces the full multi-species equilibrium under the RPM. (**Fig. 2A, SI Fig. 2A**).

**Fig. 2.**
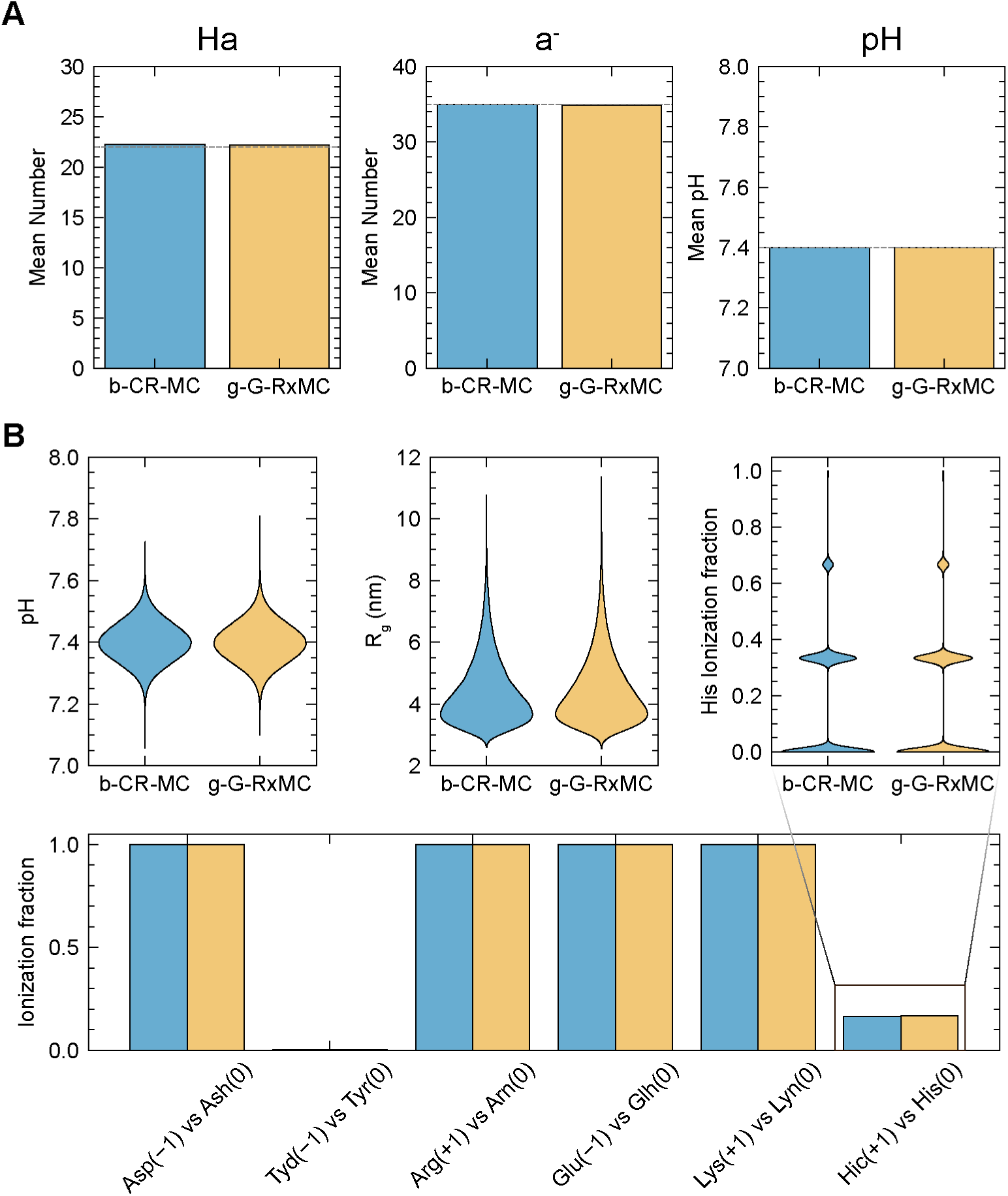
Numerical validation of the b-CR-MC algorithm against the reference g-G-RxMC method. **(A)** Ensemble-averaged buffer species populations (Ha, a⁻) and self-consistent pH in a protein-free box. Dashed lines indicate the theoretical target values. **(B)** Thermodynamic and conformational properties of a single full-length FUS (FL-FUS) chain in the dilute limit. Top panels show probability distributions (violin plots) of the self-consistent pH, radius of gyration (Rg), and Histidine ionization fraction. The bottom panel explicitly compares the mean ionization fractions across all titratable residue types. Across all panels, b-CR-MC yields statistically indistinguishable results from the reference method, confirming its accuracy.

For FL-FUS, buffer populations, self-consistent pH, and residue-resolved ionization fractions α across all six titratable types remain in quantitative agreement (**Fig. 2B, SI Fig. 2B**); as expected in the dilute limit, the net charge of a single chain constitutes a negligible perturbation to the reservoir buffer capacity. Histidine (pKa = 6.54), whose intrinsic pKa lies closest to the simulation pH and therefore exhibits the highest titration activity, provides the most stringent test near the ionization midpoint; both methods reproduce the characteristic three-peak ionization-fraction distribution within statistical uncertainty, despite FL-FUS containing only three His residues (**Fig. 2B**). Radius of gyration (Rg) distributions are statistically indistinguishable between methods (**Fig. 2B, SI Fig. 2B**), confirming that the thermodynamic coupling between ionization state and polymer conformation is preserved under the RPM approximation.

These benchmarks confirm that the reduction in reaction pool size does not compromise accuracy in either protein-free buffer systems or sequence-resolved macromolecular environments. The b-CR-MC framework therefore provides a computationally efficient yet thermodynamically equivalent foundation for condensate-scale simulations, with efficiency gains that scale favorably with sequence heterogeneity. The quantitative impact of ion-size heterogeneity on this equivalence is assessed in the Discussion.

### C. Protein Net Charge Determines the Direction of Condensate pH Gradients

A central question in condensate biophysics is whether the internal pH of a biomolecular condensate is set by the bulk reservoir or shaped by the sequence-encoded charge properties of the constituent proteins. If charge regulation is a first-order determinant of condensate electrochemistry, proteins with opposite net charges should generate pH gradients of opposite sign upon condensation.

We applied the b-CR-MC framework to dual-box phase coexistence simulations of full-length FUS (FL-FUS) and full-length PGL-3 (FL-PGL-3) (**Fig. 3**, see **Methods** for details). FUS and PGL-3 exhibit contrasting sequence compositions that govern their distinct electrochemical behaviors. At physiological pH, FUS (526 residues, 127 titratable sites) carries a net positive charge of approximately +14, driven by an Arg-rich low-complexity domain (37 Arg, 14 Lys) paired with a relatively modest acidic content (26 Asp, 11 Glu), with a calculated isoelectric point of pI = 9.84. PGL-3 (693 residues, 189 titratable sites), in contrast, bears a net negative charge of approximately -20, reflecting a substantially higher acidic load (39 Asp, 56 Glu) relative to its basic residues (44 Lys, 31 Arg), with pI = 4.25. The pI values were calculated with the same pKa values used in the simulations and excluding N- and C-terminal charges. Moreover, the sequence charge decoration (SCD) further distinguishes the two proteins.^49^ Evaluated using dilute-solution Henderson–Hasselbalch ionization fractions at pH 7.40, FUS has an SCD of 0.539, whereas PGL-3 exhibits a higher value of 0.926, reflecting the distinct charge characteristics of the two proteins.

**Fig. 3.**
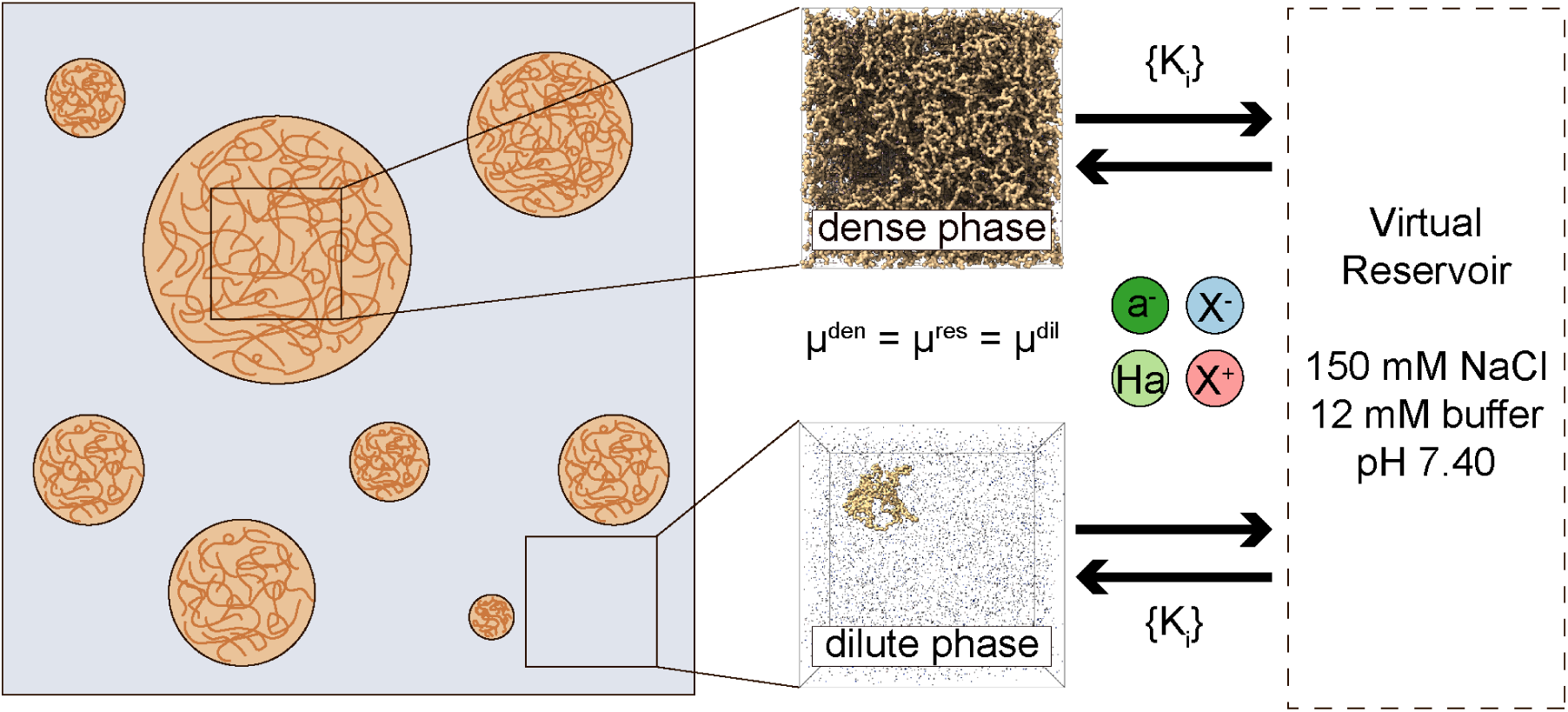
Schematic of the dual-box simulation protocol for modeling phase coexistence. A macroscopic view of liquid-liquid phase separation (left) is mapped onto two independent simulation boxes (center) representing the protein-rich dense phase and the protein-poor dilute phase. To maintain thermodynamic equilibrium without a physical interface, both phases are indirectly coupled to a shared virtual reservoir (right), which regulates the exchange of buffer (Ha, a⁻) and salt (X⁺, X⁻) ions to impose identical chemical potentials across the systems.

The simulations reveal that this sequence-encoded charge asymmetry is directly reflected in the dense-phase electrochemistry of each condensate. For FL-FUS, the effective pH increases from 7.400 ± 0.001 in the dilute phase to 7.715 ± 0.002 in the dense phase, corresponding to an alkaline shift of +0.32 pH units (**Fig. 4A, SI Fig. 3A**). FL-PGL-3 exhibits the opposite trend: the pH decreases from 7.398 ± 0.001 in the dilute phase to 7.108 ± 0.004 inside the condensate, yielding an acidic shift of −0.29 pH units (**Fig. 4B, SI Fig. 3B**). Both the sign and the sequence specificity of these simulated gradients are in qualitative agreement with experimental observations by Ausserwöger et al., confirming that condensate dense phases spontaneously shift their internal pH toward the protein isoelectric point.^3^ These results indicate that the direction of the condensate pH gradient is determined by the sign of the protein net charge relative to the reservoir pH—a sequence-encoded property that fixed-charge models cannot capture.

**Fig. 4.**
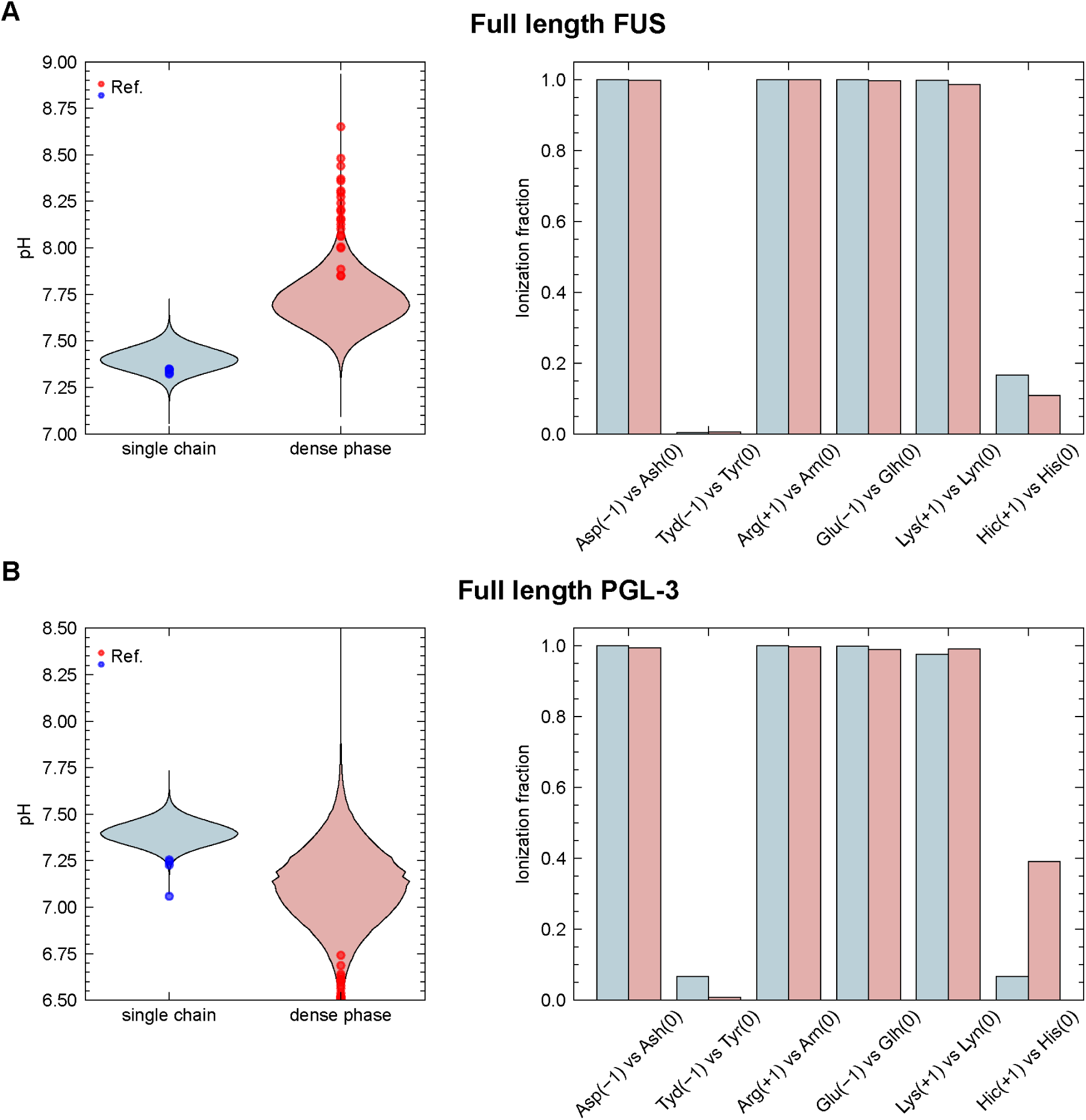
Phase-dependent charge regulation and microenvironmental pH shifts in protein condensates. **(A)** Simulation results for full-length FUS (FL-FUS). The left panel displays probability distributions (violin plots) of the effective pH for a single chain (light blue) versus the dense phase (light red), overlaid with reference pH values (solid dots) reported by Ausserwöger et al.^3^ The right panel compares the corresponding mean ionization fractions of titratable residues between the two environments. **(B)** Corresponding pH distributions and per-residue ionization fractions for full-length PGL-3 (FL-PGL-3), highlighting sequence-specific charge regulation upon condensation.^3^

### D. Charge Redistribution at the Residue Level Drives pH Shifts Toward the Isoelectric Point

These opposing pH shifts emerge from the charge-capacitance mechanism, described theoretically by Lund and Jönsson for isolated proteins^14,50^, which operates here at the condensate scale across many interacting chains. Upon entering the high-density condensate environment, protein chains experience amplified cross-chain electrostatic repulsion; the collective adjustment of titratable residue ionization fractions serves as the thermodynamic response that mitigates the resulting electrostatic free-energy penalty. Although the near-neutral simulation pH (7.40) places most titratable residues far from their intrinsic pKa values, limiting the absolute extent of charge redistribution, b-CR-MC still resolves clear shifts in ionization fractions between the two phases.

The sequence-specific proton redistribution between protein chains and their local solvent shifts the phase-local pH in opposite directions for the two proteins, establishing an electrochemical gradient across the phase boundary that is driven by charge regulation and self-consistently stabilized by the emergent Donnan potential. In the net-positive FL-FUS condensate, basic residues, particularly Lys and His, undergo partial deprotonation upon entering the dense phase, lowering the net positive charge per chain from +14.376 ± 0.006 in the single-chain limit to +14.002 ± 0.003 in the dense phase. This deprotonation releases protons into the local solvent, elevating the [a⁻]/[Ha] ratio and generating the observed alkaline shift. (**Fig. 4A** and **SI Fig. 4A**). In the net-negative FL-PGL-3 condensate, acidic residues suppress their ionization while basic residues increase their protonation, collectively shifting the net charge from −21.273 ± 0.057 to −17.037 ± 0.004, a net reduction toward charge neutrality. This collective proton uptake depresses the [a^-^]/[Ha] ratio, driving the corresponding acidic pH shift (**Fig. 4B** and **SI Fig. 4B**). In both cases, the dense phase drives the chain charge magnitude toward zero—consistent with the charge-capacitance principle that electrostatic free energy is minimized by reducing net charge in a high-density environment.^14,50^

Although the near-neutral simulation pH places most titratable residues far from their intrinsic pKa values, limiting the absolute extent of charge redistribution, b-CR-MC resolves clear, statistically significant shifts in ionization fractions between phases, indicating that even modest residue-level proton redistribution is sufficient to sustain measurable interphase pH gradients. The self-consistent Donnan potential arising from grand-canonical sampling simultaneously reinforces this asymmetry by modulating ion partitioning across the phase boundary.

### E. Sequence Composition Governs the Magnitude and Directionality of Buffer Partitioning

Beyond shifting internal pH, the two types of condensates exhibit distinct buffer partitioning patterns that reveal clear mechanistic differences in charge regulation between net-positive and net-negative systems. Phase-resolved buffer concentrations provide direct quantitative evidence of these distinct behaviors.

For FL-FUS, the neutral conjugate acid Ha is substantially depleted in the dense phase relative to the dilute phase ([Ha]dense = 2.24 mM vs. [Ha]dilute = 4.60 mM, **SI Fig. 3A**), while the charged conjugate base a⁻ remains nearly uniformly distributed across the phase boundary ([a⁻]dense = 7.17 mM vs. [a⁻] dilute = 7.27 mM, **SI Fig. 3A**). This asymmetry reflects competing thermodynamic forces: local shifting of the buffer equilibrium consumes Ha, while the near-constancy of a⁻ arises from a balance between consumption-driven depletion and the Donnan accumulation of anionic species into the net-positive condensate interior.^16,18^ Together, these effects shift the local [a⁻]/[Ha] ratio to yield the self-consistent alkaline pH gradient, resulting in a modest 21% reduction in total buffer concentration from 11.87 mM in the dilute phase to 9.41 mM in the dense phase.

In contrast, FL-PGL-3 exhibits a qualitatively distinct and more extreme pattern: both Ha and a⁻ are strongly depleted in the dense phase ([Ha]dense = 1.68 mM vs. [Ha]dilute = 4.61 mM; [a⁻]dense = 1.38 mM vs. [a⁻]dilute = 7.26 mM, **SI Fig. 3B**), with the total buffer concentration reduced by ∼74% (from 11.87 mM to 3.06 mM). Here, proton release by acidic residues consumes a-, depressing the [a⁻]/[Ha] ratio and driving the acidic pH shift; this depletion is further amplified by Donnan exclusion of the negatively charged conjugate base from the net-negative condensate interior.^16,18^

This contrast establishes that buffer partitioning is not a passive consequence of generic macromolecular crowding, but a sequence-encoded process governed by both the sign of the net protein charge and the composition of dominant titratable residues. The pronounced buffer depletion observed in PGL-3 condensates provides a molecular rationale for the experimentally reported sensitivity of condensate pH to buffer composition, suggesting that net-negative condensates of this type possess a markedly lower capacity to buffer against external pH perturbations than their net-positive counterparts.

### F. A Slab-Geometry Simulation Resolves the Spatial Structure of Condensate Electrochemistry

While the dual-box protocol employed above quantifies interphase electrochemical differences between bulk dense and dilute phases, it does not resolve how ionic species and pH vary continuously across the physical condensate interface. To probe this interfacial structure directly, we performed a complementary slab-geometry simulation of FL-FUS (**Fig. 5A**), wherein a protein-dense liquid phase forms spontaneously within a single elongated simulation cell, in direct contact with the flanking dilute regions. One-dimensional concentration profiles of Ha, a-, and monovalent salt ions (X+, X-) along the z-axis were computed from ensemble-averaged density maps, and the spatially resolved local pH was extracted at each position via the Henderson–Hasselbalch relation (**Fig. 5B**).

**Figure 5.**
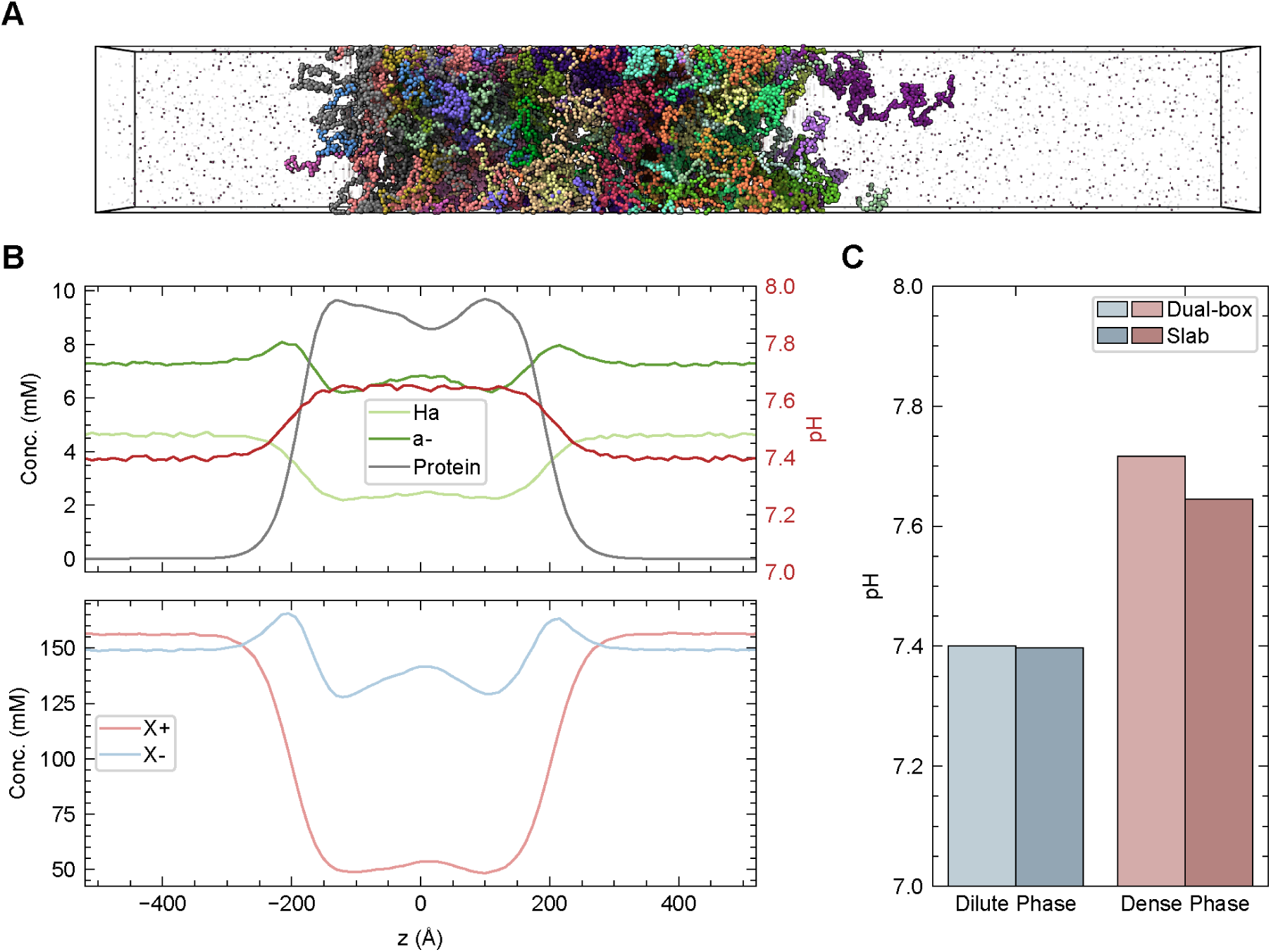
Electrochemical profiles and method validation for FUS phase coexistence.(A) Representative snapshot of the FL-FUS phase-coexistence at the end of an explicit slab-geometry simulation. **(B)** One-dimensional density and electrochemical profiles along the z-axis. (Top) Concentration profiles of the FUS protein (grey), protonated buffer (Ha, light green), and deprotonated buffer (a-, dark green) correspond to the left axis, while the spatially resolved local pH (red line) is plotted against the right axis. The condensate interior exhibits a pronounced alkaline shift. (Bottom) Concentration profiles of the monovalent salt ions reveal significant exclusion of cations (X+, light red) relative to anions (X-, light blue) from the positively charged FUS condensate, illustrating strong Donnan partitioning. **(C)** Comparison of the effective pH in the dilute (blue bars) and dense (red bars) phases computed via the dual-box protocol (lighter shades) and direct slab simulations (darker shades). Both methodologies rigorously capture the targeted macroscopic reservoir pH (7.40) in the dilute phase and robustly reproduce the alkaline shift within the condensate.

The resulting slab profiles are in qualitative agreement with the dual-box findings and provide additional spatial detail (**Fig. 5C**). Within the protein-dense slab region, the [a^-^]/[Ha] ratio is elevated relative to the flanking dilute regions, reproducing the sign and direction of the alkaline pH gradient obtained from the dual-box protocol. The local pH rises above the reservoir value of 7.4 inside the condensate, while the surrounding dilute regions are maintained near 7.4, in agreement with the single-chain reference simulations. The composite ion profiles additionally reveal a marked enrichment of anions (X^-^) relative to cations (X^+^) inside the condensate interior, a direct spatial signature of Donnan partitioning consistent with the net-positive charge of FL-FUS at this physiological pH. Because H⁺ and OH⁻ concentrations are negligible at this pH, the composite ion species primarily represent the bulk salt (NaCl), yielding profiles that closely align with findings from a previous all-atom MD study.^51^ Both the pH gradient and the ion asymmetry emerge from the same self-consistent grand-canonical sampling—without any parameterization of the interface—confirming that b-CR-MC directly resolves the continuous electrochemical structure across a condensate phase boundary within a single simulation cell.

We note, however, that the current slab simulations did not yield a discernible coexisting dilute protein phase: all 40 chains partitioned entirely into the dense slab, driving the effective dilute-phase saturation concentration toward zero. Three factors likely contribute to this outcome. First, the explicit representation of monovalent ions in b-CR-MC introduces finite excluded-volume interactions between ions and protein beads. These interactions, which are absent in standard HPS-Urry simulations that treat salt implicitly via a screened Debye–Hückel potential, may suppress chain partitioning into the dilute phase and retard chain diffusion over accessible timescales. Second, replacing the screened Debye–Hückel potential with a full particle-particle particle-mesh (PPPM) electrostatic treatment likely strengthens protein–protein interactions, further depleting the dilute phase.^42,48^ Finally, insufficient sampling cannot be ruled out: the inherently slow multi-chain dynamics of protein condensates are further compounded by the computational overhead of reactive Monte Carlo moves, making adequate conformational exploration challenging within accessible timescales. Consequently, directly extracting tie-line endpoints or saturation concentrations from the slab geometry is not feasible under the current protocol. Nevertheless, the dense-phase electrochemical observables reported here remain unaffected, as residue ionization fractions for chains occupying the dense slab are in close agreement with those from the bulk dense-phase simulation (**SI Fig. 6**). Extending slab simulations to a regime that maintains a well-defined coexisting dilute protein phase remains an objective for future work and will likely require revised ion–protein interaction parameterizations.

## IV. CONCLUSIONS AND OUTLOOK

Conventional models often approximate biomolecular condensates as electrochemically uniform; however, this is fundamentally inconsistent with recent experimental evidence demonstrating that condensates sustain spontaneous pH gradients at equilibrium.^1–3^ The results presented here provide a sequence-resolved, thermodynamically rigorous account of how these gradients arise. Rather than reflecting external perturbation or kinetic trapping, the pH shifts observed in FL-FUS (+0.32 pH units, alkaline) and FL-PGL-3 (−0.29 pH units, acidic) emerge self-consistently from grand-canonical charge regulation. The dense-phase protein environment shifts residue ionization fractions toward net charge neutrality, redistributing protons between the protein chains and the local solvent in a direction and magnitude determined by the protein’s net charge relative to the reservoir pH. This charge-capacitance mechanism, described theoretically by Lund and Jönsson for isolated proteins,^14,50^ operates here at the condensate scale across many interacting chains. It is therefore a directly sequence-encoded, emergent property inaccessible to fixed-charge frameworks.

The contrasting behaviors of FL-FUS and FL-PGL-3 demonstrate that the sign of net charge is the primary sequence determinant of pH gradient directionality. Both proteins phase-separate under the same reservoir conditions (150 mM NaCl, 12 mM buffer, pH 7.40), yet they generate pH gradients of opposite sign because their isoelectric points bracket the reservoir pH from above (FUS, pI = 9.84) and below (PGL-3, pI = 4.25). The qualitative agreement between these simulated gradients and the experimental observations of Ausserwöger et al. supports the charge-regulation interpretation and establishes b-CR-MC as a sequence-resolved predictive platform for electrochemical characterization across sequence space^3^. The modest quantitative attenuation of the computed pH shifts relative to experimental values is likely primarily driven by two well-defined model approximations. First, substituting the experimental buffer with a monovalent surrogate alters Donnan-driven exclusion and omits specific multivalent interactions or intra-condensate accumulation.^32^ Second, the omission of RNA or other polyanionic clients eliminates an additional biological driver of electrochemical asymmetry.^3^ Both limitations are tractable and do not alter the mechanistic conclusions.

The internal alkaline pH shift within FL-FUS condensates, predicted here as +0.32 pH units and experimentally verified at ∼+1.0 pH units^3^, shifts the dense-phase interior toward the protein’s isoelectric point, thereby reducing net interchain electrostatic repulsion, which might play a role of promoting the aging of droplets and liquid-to-solid transition.^52,53^ Because the pKa values of His (6.54), Lys (10.53), and Arg (12.48) span a wide range, this dense-phase-specific alkaline shift preferentially deprotonates histidine residues, altering FUS charge patterning in a residue-position-dependent manner that remains invisible to conventional fixed-charge models. This shifting electrochemical landscape intrinsically modulates the biological readout of post-translational modifications (PTMs), such as serine/tyrosine phosphorylation and arginine methylation, and determines how charge-altering, ALS-associated missense mutations in the low-complexity domain may alter internal pH to amplify or dampen the charge redistribution mechanism.^54–60^ Beyond homotypic stress granule dynamics, this electrochemical coupling extends to the DNA damage response, where FUS phase-separates at DNA double-strand breaks to recruit polyanionic clients such as nucleic acids;^5,61–63^ recruitment of these clients may drive secondary, composition-dependent pH shifts,^3^ that might dynamically modulate the electrostatic interaction networks of recruited DNA repair factors. By treating the reciprocal coupling among macromolecular composition, differential ion partitioning, and titratable charge states on an equal thermodynamic footing, b-CR-MC provides a sequence-to-electrochemistry map that can quantify how both disease-variant sequences and multicomponent client composition alter local electrostatic environments, with direct implications for DNA repair efficiency and disease progression.

The present study establishes the mechanistic framework and validates the computational tool; however, several methodological and biological extensions are needed to bridge the gap between these CG predictions and quantitative experimental measurements. The monovalent buffer representation employed here, while necessary for RPM compatibility, alters Donnan-driven exclusion and omits potential specific multivalent interactions and intra-condensate accumulation.^32^ Extending b-CR-MC to multivalent buffer species via the full g-G-RxMC framework is therefore a priority. Additionally, ion-size heterogeneity might introduce a slight pH unit error in dilute-phase ionization that may be amplified by macromolecular crowding in the dense phase; a systematic benchmarking of this error against all-atom or ion-specific CG models is warranted. Although the current slab-geometry implementation is limited by explicit-ion excluded-volume interactions that suppress dilute-phase chain partitioning, preventing the direct extraction of tie-line endpoints, the geometry itself remains the most direct route to spatially resolved interfacial electrochemical profiles. Future efforts combining improved ion-protein force-field parameterizations with extended simulation timescales and Gibbs-ensemble coupling methods would enable the simultaneous, self-consistent determination of both coexistence concentrations and phase-resolved electrochemistry within a unified simulation framework. Beyond these technical extensions, the b-CR-MC framework is well positioned to advance sequence-level investigations into how mutations, post-translational modifications, and multi-component compositions shape condensate electrochemistry, thereby opening a direct computational route to several outstanding biological questions: how do multi-component condensates, containing both positively and negatively charged protein clients alongside RNA, modulate internal pH relative to single-component systems? Does the electrochemical gradient across the condensate interface generate an electric field sufficient to influence the orientation or partitioning of dipolar client molecules? Can sequence design principles derived from the charge-capacitance mechanism be used to engineer condensates with prescribed internal pH values, enabling synthetic biology applications in which condensate electrochemistry is a tunable design parameter? These questions are now computationally tractable with the b-CR-MC framework, which provides a thermodynamically exact, sequence-resolved foundation for connecting protein sequence, condensate composition, and emergent electrochemical function.

## SUPPLEMENTARY MATERIAL

See the supplementary material for the complete reaction sets of the original g-G-RxMC framework and the mathematical proof of thermodynamic equivalence between the b-CR-MC and g-G-RxMC frameworks; the derivation and validation of the virtual-particle strategy for grand-canonical sampling; benchmarking results for the custom HOOMD-blue plugin; and additional simulation details including explicit protein sequences, interaction parameters, phase-resolved buffer distributions, and residue-level ionization fractions.

## Supporting information

Supplementary Information containing additional text, figures, and tables supporting the main manuscript.

## ACKNOWLEDGMENTS

This work was supported by the National Institute of General Medical Sciences of the National Institutes of Health under grant R35GM153388. Y.C.K. acknowledges support from the U.S. Naval Research Laboratory through the Office of Naval Research Base Program. We thank Dr. Qizan Chen for assistance in improving the HOOMD-blue plugin. We also gratefully acknowledge the Texas A&M High Performance Research Computing facility for providing the computational resources essential to the simulations reported in this study.

## AUTHOR DECLARATIONS

### Conflict of Interest

The authors declare no conflict of interest.

## DATA AVAILABILITY

The data that support the findings of this study are available within the article and its supplementary material. The source code for the custom HOOMD-blue plugin developed in this work is openly available on GitHub at https://github.com/slweng0321/RxMC.git.

## Notes

### Competing Interest Statement

The authors have declared no competing interest.

