## Supplementary Information containing additional text, figures, and tables supporting the main manuscript. for "Molecular Origins of pH Gradients in Charge-Regulated Biomolecular Condensates"

### Supplementary Material for: Molecular Origins of pH Gradients in Charge-Regulated Biomolecular Condensates

#### Section S1. THEORETICAL FRAMEWORK: FROM g-G-RxMC TO b-CR-MC

This section provides the complete reaction sets for G-RxMC and g-G-RxMC, together with the proof of thermodynamic equivalence between b-CR-MC and g-G-RxMC under the Restricted Primitive Model (RPM), as referenced in the main text Materials and Methods.

##### A. Grand-Reaction Monte Carlo (G-RxMC)

The Grand-Reaction Monte Carlo (G-RxMC) method, introduced by Landsgesell et al., simulates the acid–base equilibria between weak polyelectrolytes and a macroscopic reservoir of fixed composition.[1] The method couples explicit acid–base reactions within the simulation box to grand-canonical insertion and deletion of ion pairs from an implicit reservoir, tracking each ionic species ($Na^+$, $Cl^-$, $H^+$, $OH^-$) individually. The reservoir is characterized by a set of equilibrium constants $\{K_i\}$ that encode the chemical potentials of all exchangeable species.

For a system at fixed reservoir pH in the presence of NaCl at concentration $c_{NaCl}$, four ion-pair exchange reactions are required:

$$\emptyset \rightleftharpoons \mathrm{Na}^+ + \mathrm{Cl}^- \quad (K_{\mathrm{NaCl}})$$

$$\emptyset \rightleftharpoons \mathrm{H}^+ + \mathrm{Cl}^- \quad (K_{\mathrm{HCl}})$$

$$\emptyset \rightleftharpoons \mathrm{Na}^+ + \mathrm{OH}^- \quad (K_{\mathrm{NaOH}})$$

$$\emptyset \rightleftharpoons \mathrm{H}^+ + \mathrm{OH}^- \quad (K_w)$$

(In our HOOMD-blue implementation, the null state (∅) is represented computationally by an equivalent number of virtual particles, the role of which is detailed in **Section S2**.)

These reactions are linked by the thermodynamic constraint

$$K_{\mathrm{NaCl}} \cdot K_w = K_{\mathrm{HCl}} \cdot K_{\mathrm{NaOH}}$$

Each titratable residue type contributes to four dissociation reactions and one identity-swap reaction:

$$\mathrm{HA} \rightleftharpoons \mathrm{H}^+ + \mathrm{A}^- \quad (K_a)$$

$$\mathrm{HA} + \mathrm{OH}^- \rightleftharpoons \mathrm{A}^- \quad (K_{\mathrm{HA,OH^-}})$$

$$\mathrm{HA} \rightleftharpoons \mathrm{Na}^+ + \mathrm{A}^- \quad (K_{HA,\mathrm{Na^+}})$$

$$\mathrm{HA} + \mathrm{Cl}^- \rightleftharpoons \mathrm{A}^- \quad (K_{HA,\mathrm{Cl^-}})$$

$$\mathrm{HA}_i + \mathrm{A}_j^- \rightleftharpoons \mathrm{A}_i^- + \mathrm{HA}_j \quad (K_{swap} = 1)$$

##### B. Generalized Formulation for Buffer Exchange (g-G-RxMC)

Beyer and Holm generalized the G-RxMC framework to allow the exchange of weak polyprotic buffer acid species between the simulation box and the reservoir.[2] In this generalized scheme (g-G-RxMC), the reservoir pH is not an external input but emerges self-consistently from the acid–base equilibrium of the buffer at a specified total concentration. For a monovalent buffer acid Ha with dissociation constant $K_{bf}$, the following six additional reactions enter the pool:

$$\emptyset \rightleftharpoons \mathrm{Ha} \quad (K_{\mathrm{Ha}})$$

$$\emptyset \rightleftharpoons \mathrm{a}^- + \mathrm{Na}^+ \quad (K_{\mathrm{a}^-,\mathrm{Na}^+})$$

$$\mathrm{Ha} \rightleftharpoons \mathrm{a}^- + \mathrm{H}^+ \quad (K_{\mathrm{bf}})$$

$$\mathrm{Ha} + \mathrm{OH}^- \rightleftharpoons \mathrm{a}^- \quad (K_{\mathrm{bf}}/K_w)$$

$$\mathrm{Ha} \rightleftharpoons \mathrm{a}^- + \mathrm{Na}^+ \quad (K_{\mathrm{bf,Na}} = K_{\mathrm{bf}} \cdot K_{\mathrm{NaCl}}/K_{\mathrm{HCl}})$$

$$\mathrm{Ha} + \mathrm{Cl}^- \rightleftharpoons \mathrm{a}^- \quad (K_{\mathrm{bf}}/K_{\mathrm{HCl}})$$

The reaction $\emptyset \rightleftharpoons \mathrm{H}^+ + \mathrm{a}^-$ $(K_{\mathrm{H}^+,\mathrm{a}^-})$, formally required for completeness in the g-G-RxMC framework of Beyer and Holm for n = 1, is omitted from our sampling pool.[2] This omission does not compromise thermodynamic rigor, as the reaction is thermodynamically redundant: exchange of these species is fully captured by the sequential neutral-acid insertion ($\emptyset \rightleftharpoons \mathrm{Ha}$) followed by the dissociation reaction ($\mathrm{Ha} \rightleftharpoons \mathrm{a}^- + \mathrm{H}^+$), both of which are explicitly included. Any sufficient subset of moves that satisfies the stoichiometric rank condition converges to the same thermodynamically exact equilibrium state. Moreover, at the target pH of 7.4, the concentration of free $H^+$ ions is vanishingly small, rendering the acceptance probability for a direct pair-insertion move negligible in practice. As established in prior work, redundant reaction sets may enhance sampling efficiency in certain regimes but do not alter equilibrium observables.[1] For a monovalent buffer system containing N types of titratable residues, the g-G-RxMC reaction pool used in the present work comprises 10 + 5 N distinct reactions, corresponding to the full 11 + 5N pool of Beyer and Holm with the thermodynamically redundant $H^+$/$a^-$ pair-insertion channel excluded.

##### C. Proof of Thermodynamic Equivalence Between b-CR-MC and g-G-RxMC

The b-CR-MC framework introduced in the main text reduces the g-G-RxMC reaction pool by merging monovalent ions under the RPM. The following proof establishes that this merging is thermodynamically exact: the grand-canonical partition functions of b-CR-MC and g-G-RxMC are identical under the RPM, and consequently all thermodynamic observables are equivalent in both formulations.

Consider a system at fixed volume V and temperature T coupled to a reservoir fixing the chemical potentials $\mu_{H^+}$, $\mu_{Na^+}$, $\mu_{OH^-}$, $\mu_{Cl^-}$, $\mu_{Ha}$, and $\mu_{a^-}$. The g-G-RxMC grand-canonical partition function is:

$$\Xi_{\text{full}} = \sum_{\{N_s\}} \frac{e^{\beta \sum_s \mu_s N_s}}{N_{H^+}!\, N_{Na^+}!\, N_{OH^-}!\, N_{Cl^-}!\, N_{Ha}!\, N_{a^-}!} \int d\boldsymbol{r}^N\, e^{-\beta U(\boldsymbol{r})}$$

where the sum runs over all combinations of non-negative integer particle numbers $\{N_s\}$ for each species, and U(**r**) is the potential energy of the system.

Under the RPM, any microstate, containing $N_{H^+}$ protons and $N_{Na^+}$ sodium ions, is physically indistinguishable from a state obtained by relabeling individual cations. The Boltzmann weight therefore depends only on the total cation count $N_+ = N_{H^+} + N_{Na^+}$ and total anion count $N_- = N_{OH^-} + N_{Cl^-}$. Summing over all cation labelings at fixed $N_+$ yields:

$$\sum_{N_{H^+}=0}^{N_+} \frac{e^{\beta(\mu_{H^+} N_{H^+} + \mu_{Na^+}(N_+ - N_{H^+}))}}{N_{H^+}!\, (N_+ - N_{H^+})!} = \frac{(\lambda_{H^+} + \lambda_{Na^+})^{N_+}}{N_+!}$$

where $\lambda_s \equiv e^{\beta \mu_s}$ is the fugacity of species $s$. Defining the merged fugacity $\lambda_{X^+} \equiv \lambda_{H^+} + \lambda_{Na^+}$ and $\lambda_{X^-} \equiv \lambda_{OH^-} + \lambda_{Cl^-}$, with the corresponding merged chemical potential:

$$\mu_{X^+} \equiv k_B T \ln(\lambda_{H^+} + \lambda_{Na^+}) = k_B T \ln\left(e^{\beta \mu_{H^+}} + e^{\beta \mu_{Na^+}}\right)$$

$$\mu_{X^-} \equiv k_B T \ln(\lambda_{OH^-} + \lambda_{Cl^-}) = k_B T \ln\left(e^{\beta \mu_{OH^-}} + e^{\beta \mu_{Cl^-}}\right)$$

the partition function simplifies to:

$$\Xi_{\text{full}} = \sum_{N_+, N_-, N_{Ha}, N_{a^-}} \frac{e^{\beta(\mu_{X^+} N_+ + \mu_{X^-} N_- + \mu_{Ha} N_{Ha} + \mu_{a^-} N_{a^-})}}{N_+!\, N_-!\, N_{Ha}!\, N_{a^-}!} \int d\boldsymbol{r}^N\, e^{-\beta U(\boldsymbol{r}^N)} \equiv \Xi_{\text{b-CR-MC}}$$

This expression is precisely the grand-canonical partition function of b-CR-MC, in which only the merged species $X^+$ and $X^-$, together with the buffer species Ha and $a^-$, appear explicitly. Because the two partition functions are identical, all thermodynamic observables, including mean particle numbers, charge fluctuations, free energies, and ionization fractions, are the same in both formulations. The b-CR-MC equilibrium constants follow directly from the merged fugacities:

$$K_{salt} \propto \lambda_{X^+}\lambda_{X^-}, \qquad K_{bf} \cdot \Phi \propto \lambda_{a^-}\lambda_{X^+}/\lambda_{Ha}$$

where $\Phi = 1 + K_{NaCl}/K_{HCl}$ encodes the sodium-to-proton fugacity ratio. This completes the proof that b-CR-MC is thermodynamically exact under the RPM, with no approximation beyond the indistinguishability of same-valency ions.

#### Section S2. VIRTUAL-PARTICLE STRATEGY: GRAND-CANONICAL LIMIT AND ACCEPTANCE PROBABILITY

This section provides (1) the proof that the finite virtual-particle pool converges to the true grand-canonical limit, and (2) the derivation that the virtual-particle reaction scheme preserves the correct reactive ensemble acceptance probability, as referenced in the main text Materials and Methods.

##### A. Convergence to the Grand-Canonical Limit

Because the total particle count (real plus virtual) is fixed in HOOMD-blue, the system formally occupies a finite-pool grand-canonical ensemble with partition function:

$$\mathrm{Z}(N_{\mathrm{virtual}}, \mu, V, T) = \sum_{N_1=0}^{N_1^{\max}} \cdots \sum_{N_m=0}^{N_m^{\max}} e^{\mu N} Q(N, V, T)$$

where the upper summation limit $N_j^{\max}$ for each ionic species $j$ is set by the total number of available virtual particles. This differs from the true grand-canonical partition function $\Xi(\mu, V, T)$ solely in the truncation of states involving particle numbers far larger than the mean. Because the Boltzmann weight $e^{\mu N} Q(N, V, T)$ decays exponentially for particle numbers well above the ensemble mean $\langle N_j \rangle$, truncated states are negligible provided the pool is sufficiently large, and $Z \to \Xi$ as $N_{\mathrm{virtual}} \to \infty$. In all simulations reported here, the virtual pool was sized conservatively and its adequacy was confirmed a posteriori: all species converged to stable ensemble-averaged populations, and no insertion move was rejected due to pool depletion, confirming that every simulation operated in the grand-canonical limit (**SI Fig. 5**).

##### B. Derivation of Reactive Ensemble Acceptance Probability with Virtual Particles

Here we show that employing virtual particles in HOOMD-blue to realize the grand-canonical limit preserves the correct reactive ensemble acceptance probability. Consider a grand-canonical ensemble for a system of volume V and temperature T in contact with a reservoir for four particle species {A, B, C, V} with fixed chemical potentials {$\mu_A$, $\mu_B$, $\mu_C$, $\mu_V$}, undergoing the reaction

$$\mathrm{A} + \mathrm{V} \rightleftharpoons \mathrm{B} + \mathrm{C}$$

where V denotes a virtual (non-interacting) particle. The total particle number N = $N_A$ + $N_B$ + $N_C$ + $N_V$ is fixed in HOOMD-blue, so every reaction event conserves N: one A and one V are consumed while one B and one C are produced.

For fixed particle number $\{N_A, N_B, N_C, N_V\}$, the canonical partition function is

$$Q(\{N_s\}, V, T) = \frac{1}{N_A!\, N_B!\, N_C!\, N_V!} \cdot \frac{1}{\Lambda_A^{3N_A} \Lambda_B^{3N_B} \Lambda_C^{3N_C} \Lambda_V^{3N_V}} \int dr^{N_{\text{real}}}\, e^{-\beta U(r)}$$

Because virtual particles are non-interacting and thermodynamically ideal, they do not appear in the potential energy U(**r**); only the $N_{real}$ = $N_A$ + $N_B$ + $N_C$ real-particle coordinates enter the configuration integral.

The grand-canonical partition function then becomes

$$\mathcal{Z} = \sum_{\{N_s\}} \prod_{s \in \{A,B,C,V\}} \frac{e^{\beta \mu_s N_s}}{\Lambda_s^{3N_s}\, N_s!} \cdot \int dr^{N_{\text{real}}}\, e^{-\beta U(r)}$$

and the equilibrium probability of a microstate $(\{N_s\}, r)$ is

$$\pi(\{N_s\}, r) = \frac{1}{\mathcal{Z}} \cdot \prod_{s \in \{A,B,C,V\}} \frac{e^{\beta \mu_s N_s}}{\Lambda_s^{3N_s}\, N_s!} \cdot e^{-\beta U(r)}$$

For the forward reaction move, the old state is $(\{N_A, N_B, N_C, N_V\}, r)$ and the new state is $(\{N_A - 1, N_B + 1, N_C + 1, N_V - 1\}, r')$. Detailed balance requires

$$\pi(\text{old}) \cdot P_{\text{acc}}(\text{old} \to \text{new}) = \pi(\text{new}) \cdot P_{\text{acc}}(\text{new} \to \text{old})$$

Reactive particles are chosen uniformly at random among all eligible particles; if no suitable particle is available, the move is rejected. Assuming symmetric proposal probabilities, the Metropolis acceptance criterion gives

$$P_{\text{acc}}(\text{old} \to \text{new}) = \min\left[1, \frac{\pi(\text{new})}{\pi(\text{old})}\right]$$

Substituting the microstate probabilities and collecting terms for each species gives:

$$\frac{\pi(\text{new})}{\pi(\text{old})} = \frac{N_A N_V}{(N_B + 1)(N_C + 1)} \frac{\Lambda_A^3 \Lambda_V^3}{\Lambda_B^3 \Lambda_C^3} e^{\beta(\mu_B + \mu_C - \mu_A - \mu_V)} e^{-\beta \Delta U}$$

where $\Delta U = U(\boldsymbol{r}') - U(\boldsymbol{r})$.

Each species carries a chemical potential

$$\mu_s = k_B T \ln\left(\frac{N_s}{V}\Lambda_s^3\right) + \mu_s^{\text{ex}}$$

For the virtual particle, $\mu_V^{\text{ex}} = 0$ by construction. Combining the virtual-particle factors from the chemical potential term:

$$e^{-\beta\mu_V} \cdot \Lambda_V^3 \cdot N_V = \frac{V}{N_V \Lambda_V^3} \cdot \Lambda_V^3 \cdot N_V = V$$

Both $N_V$ and $\Lambda_V$ cancel exactly, and the ideal contribution of the virtual particle reduces to a plain factor of V. Similarly, the ideal parts of $\mu_A$, $\mu_B$, $\mu_C$ combine with their thermal-wavelength factors to give $N_B N_C / (N_A V)$, with all $\Lambda_s$ cancelling. The probability ratio therefore simplifies to

$$\frac{\pi(\text{new})}{\pi(\text{old})} = \frac{N_A V}{(N_B+1)(N_C+1)} e^{\beta\Delta\mu^{ex}} e^{-\beta\Delta U}$$

where $\Delta\mu^{\text{ex}} = \mu_B^{\text{ex}} + \mu_C^{\text{ex}} - \mu_A^{\text{ex}}$. Neither $N_V$, $\Lambda_V$, nor any ideal term associated with the virtual species appears in the final expression.

The effective reaction constant $K_r$ absorbs the excess chemical-potential correction:

$$K_r \equiv K \cdot e^{\beta\Delta\mu^{\text{ex}}}, \qquad K = \frac{[B]^{\text{res}}[C]^{\text{res}}}{[A]^{\text{res}} \cdot c^\circ}$$

where $c^\circ = 1$ M. Substituting $N_s$ = [s] $N_{Avo}$ V at reservoir equilibrium into the probability ratio gives:

$$\frac{\pi(\text{new})}{\pi(\text{old})} = K_r \cdot (Vc^\circ N_{Avo})^{\Delta\nu_r} \cdot \frac{N_A}{(N_B+1)(N_C+1)} \cdot e^{-\beta\Delta U}$$

where $\Delta\nu_r = +1$ is the net change in the real particle number for the reaction $A \rightarrow B + C$. The Metropolis acceptance probability is therefore

$$P_{\text{acc}} = \min\left[1,\ K_r \cdot \left(Vc^\circ N_A^{\text{Avo}}\right)^{\Delta\nu_r} \cdot \frac{N_A}{(N_B+1)(N_C+1)} \cdot e^{-\beta\Delta U}\right]$$

This result is identical to the standard reactive Monte Carlo acceptance probability derived without virtual particles, confirming that the virtual-particle strategy introduces no bias into the equilibrium sampling. The μ-tuning procedure absorbs the excess chemical-potential offset $e^{\beta\Delta\mu^{\text{ex}}}$ into $K_r$, and the $N_V$ and $\Lambda_V$ factors cancel exactly through the ideal-gas term of the virtual particle's chemical potential. Crucially, this absorption eliminates the erroneous symmetry-breaking artifacts that can arise when using explicit ions.[3]

**Section S3. VALIDATION OF THE CUSTOM HOOMD-BLUE PLUGIN**

This section reports the benchmark simulations used to validate the custom HOOMD-blue constant-pH and reaction-ensemble plugin, as referenced in the main text Materials and Methods.

To validate the correct operation of the custom constant-pH and reaction-ensemble plugin and to assess the accuracy of the virtual-particle strategy, we performed constant-pH and reaction-ensemble simulations of weak-acid dissociation using the HOOMD-blue package (version 5.1.1).[4] The results were compared with those obtained using the ESPResSo molecular dynamics package (version 5.0.0).[5,6] The simulation setup used in both approaches followed exactly that reported in the reference study.[7] Results obtained with HOOMD-blue and ESPResSo agree with the reference data to within statistical uncertainty (**SI Fig. 1A**). As an independent cross-validation, the reaction-ensemble module was used to reproduce the CR-MC results of Curk et al. for an ideal monoacid, a weak monoacid, and a weak polyacid system.[1,8] Results from HOOMD-blue and LAMMPS (29 August 2024, Update 1) are likewise in agreement with the reference data to within statistical uncertainty (**SI Fig. 1B**).[9]

**Table S-I. Simulated sequences**

| Name | Sequence |
|---|---|
| FUS | MASNDYTQQATQSYGAYPTQPGQGYSQQSSQPYGQQSYSGYSQSTDTSGYGQSS<br>YSSYGQSQNTGYGTQSTPQGYGSTGGYGSSQSSQSSYGQQSSYPGYGQQPAPSS<br>TSGSYGSSSQSSSYGQPQSGSYSQQPSYGGQQQSYGQQQSYNPPQGYGQQNQY<br>NSSSGGGGGGGGGGNYGQDQSSMSSGGGSGGGYGNQDQSGGGGSGGYGQQD<br>RGGRGRGGSGGGGGGGGGGYNRSSGGYEPRGRGGGRGGRGGMGGSDRGGFN<br>KFGGPRDQGSRHDSEQDNSDNNTIFVQGLGENVTIESVADYFKQIGIIKTNKKTGQP<br>MINLYTDRETGKLKGEATVSFDDPPSAKAAIDWFDGKEFSGNPIKVSFATRRADFNR<br>GGGNGRGGRGRGGPMGRGGYGGGGSGGGGRGGFPSGGGGGGGQQRAGDWK<br>CPNPTCENMNFSWRNECNQCKAPKPDGPGGGPGGSHMGGNYGDDRRGGRGGY<br>DRGGYRGRGGDRGGFRGGRGGGDRGGFGPGKMDSRGEHRQDRRERPY |
| PGL-3 | MEANKRQIVEVDGIKSYFFPHLAHYLASNDELLVNNIAQANKLAAFVLGATDKRPSNE<br>EIAEMILPNDSSAYVLAAGMDVCLILGDDFRPKFDSGAEKLSQLGQAHDLAPIIDDEKK<br>ISMLARKTKLKKSNDAKILQVLLKVLGAEEAEEKFVELSELSSALDLDFDVYVLAKLLG<br>FASEELQEEIEIIRDNVTDAFEACKPLLKKLMIEGPKIDSVDPFTQLLLTPQEESIEKAVS<br>HIVARFEEASAVEDDESLVLKSQLGYQLIFLVVRSLADGKRDASRTIQSLMPSSVRAE<br>VFPGLQRSVFKSAVFLASHIIQVFLGSMKSFEDWAFVGLAEDLESTWRRRAIAELLKK<br>FRISVLEQCFSQPIPLLPQSELNNETVIENVNNALQFALWITEFYGSESEKKSLNQLQF<br>LSPKSKNLLVDSFKKFAQGLDSKDHVNRIIESLEKSSSSEPSATAKQTTTSNGPTTVST<br>AAQVVTVEKMPFSRQTIPCEGTDLANVLNSAKIIGESVTVAAHDVIPEKLNAEKNDNT<br>PSTASPVQFSSDGWDSPTKSVALPPKISTLEEEQEEDTTITKVSPQPQERTGTAWGS<br>GDATPVPLATPVNEYKVSGFGAAPVASGFGQFASSNGTSGRGSYGGGRGGDRGGR<br>GAYGGDRGRGGSGDGSRGYRGGDRGGRGSYGEGSRGYQGGRAGFFGGSRGGS |

**Table S-II. Parameters for titratable residues and ion species**

| Amino acid | ε | σ | λ | pKa | Charge |
|---|---|---|---|---|---|
| Asp | 0.2 | 5.58 | 0.294119 | 3.67 | -q* |
| Ash | | | 0.602942 | | 0 |
| Tyr | | 6.46 | 0.897059 | 9.84 | 0 |
| Tyd | | | 0.382354 | | -q |
| Arg | | 6.56 | 0.558824 | 12.48 | +q |
| Arn | | | 0.958824 | | 0 |
| Glu | | 5.92 | 0.000000 | 4.25 | -q |
| Glh | | | 0.647060 | | 0 |
| Lys | | 6.36 | 0.382354 | 10.53 | +q |
| Lyn | | | 0.632354 | | 0 |
| His | | 6.08 | 0.764707 | 6.54 | 0 |
| Hic | | | 0.647060 | | +q |
| $X^+$ | 1.0 | 3.55 | 0** | n/a | +q |
| $X^-$ | | 3.55 | | n/a | -q |
| Ha | | 3.55 | | 7.20 | 0 |
| $a^-$ | | 3.55 | | | -q |

*: unit charge is scaled to 2.037

**: pure WCA repulsive interactions

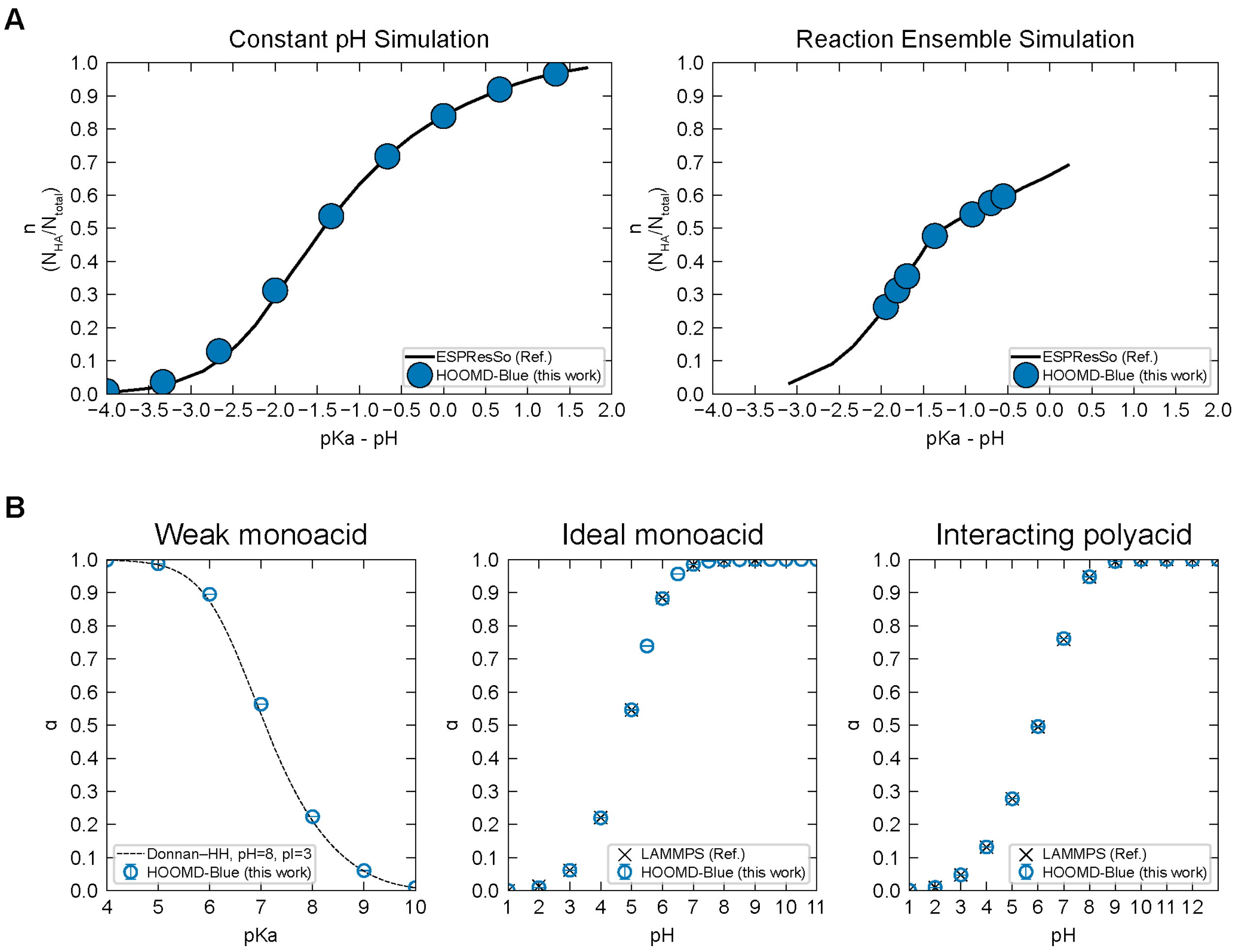


**Fig. S1. Validation of the custom HOOMD-blue constant-pH and reaction-ensemble plugins. (A)** Fractional dissociation of a weak acid in constant-pH (left) and reaction-ensemble (right) simulations. Results from the custom HOOMD-blue plugin (solid blue circles) are in quantitative agreement with reference data from the ESPResSo package (solid black lines). The simulation setup follows that reported in the reference study.[7] **(B)** Degree of dissociation (α) for a weak monoacid (left), an ideal monoacid (middle), and an interacting polyacid (right). HOOMD-blue results (open blue circles) agree with the Donnan–Henderson–Hasselbalch theoretical curve (Donnan–HH, dashed line) and LAMMPS reference simulations (black crosses) to within statistical uncertainty.[8]

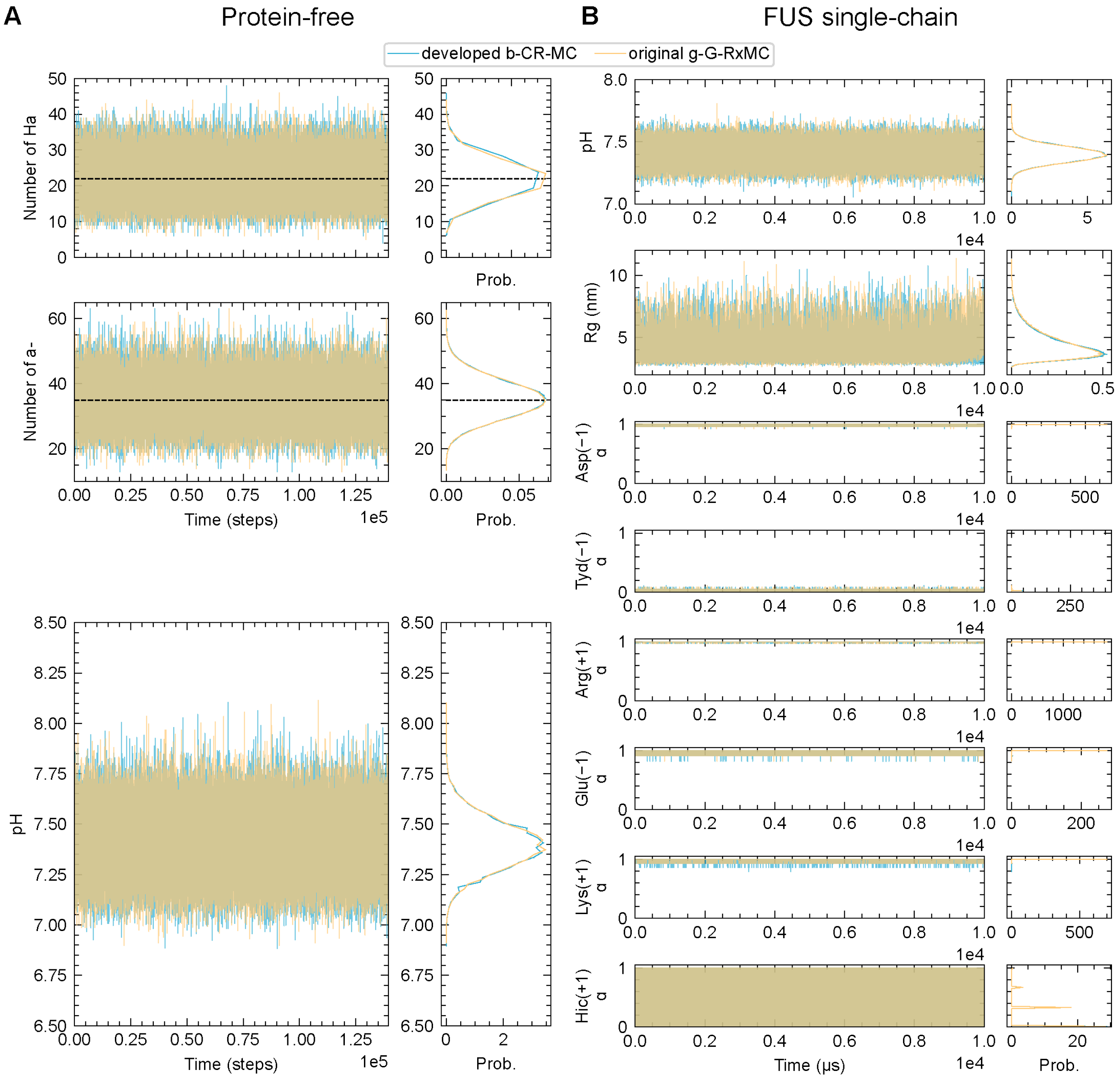


**Fig. S2. Time-series validation of b-CR-MC against g-G-RxMC. (A)** Time-series trajectories (left) and corresponding probability distributions (right) for a protein-free box. The panels track the instantaneous fluctuations of the buffer species (Ha and $a^-$) and the effective pH. Dashed black lines indicate the target values. **(B)** Conformational and charge dynamics for a single full-length FUS chain, comparing the time evolution and probability densities of the local pH, the radius of gyration (Rg), and the ionization fractions (α) for all titratable residue types (Asp, Tyr, Arg, Glu, Lys, His). The close overlap between the two methods confirms the quantitative equivalence of the b-CR-MC and g-G-RxMC frameworks.

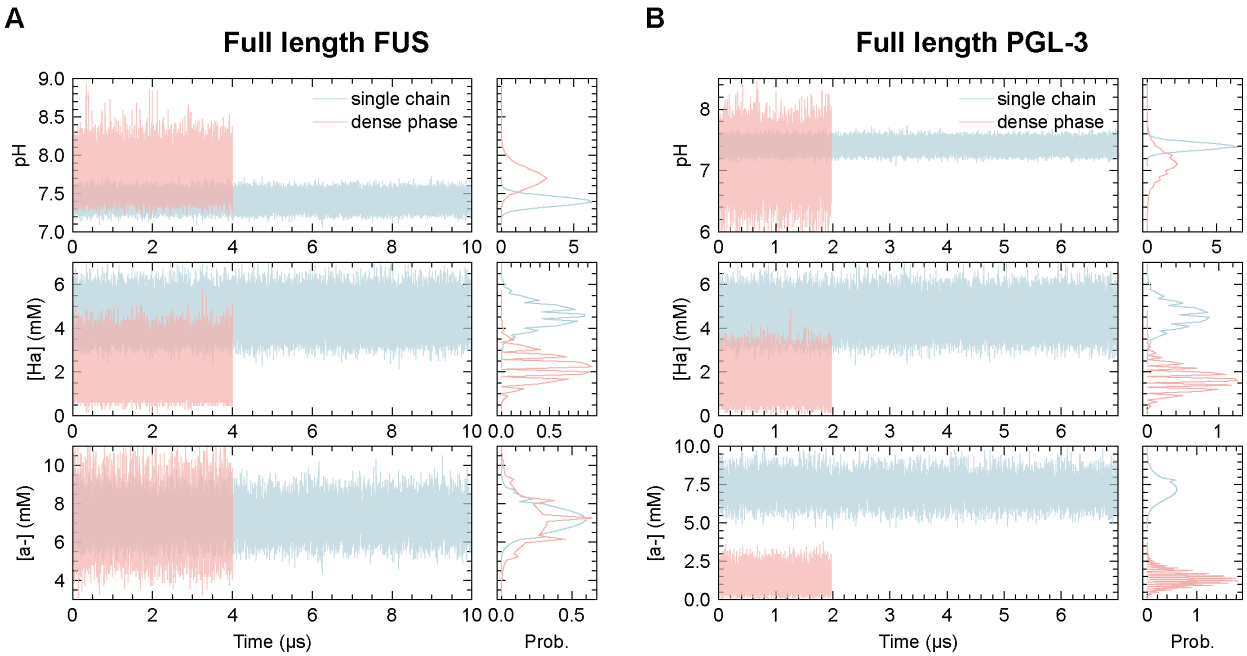


**Fig. S3. Phase-resolved electrochemical microenvironments of FL-FUS and FL-PGL-3.** Simulation trajectories (left subpanels) and corresponding probability density distributions (right subpanels) comparing a single isolated chain (dilute-phase proxy, light blue) with the dense phase of the dual-box coexistence simulation (pink). Data are shown for **(A)** full-length FUS and **(B)** full-length PGL-3. Each panel reports the time evolution of the effective local pH and the local concentrations of the neutral ([Ha]) and anionic ([a$^-$]) buffer species.

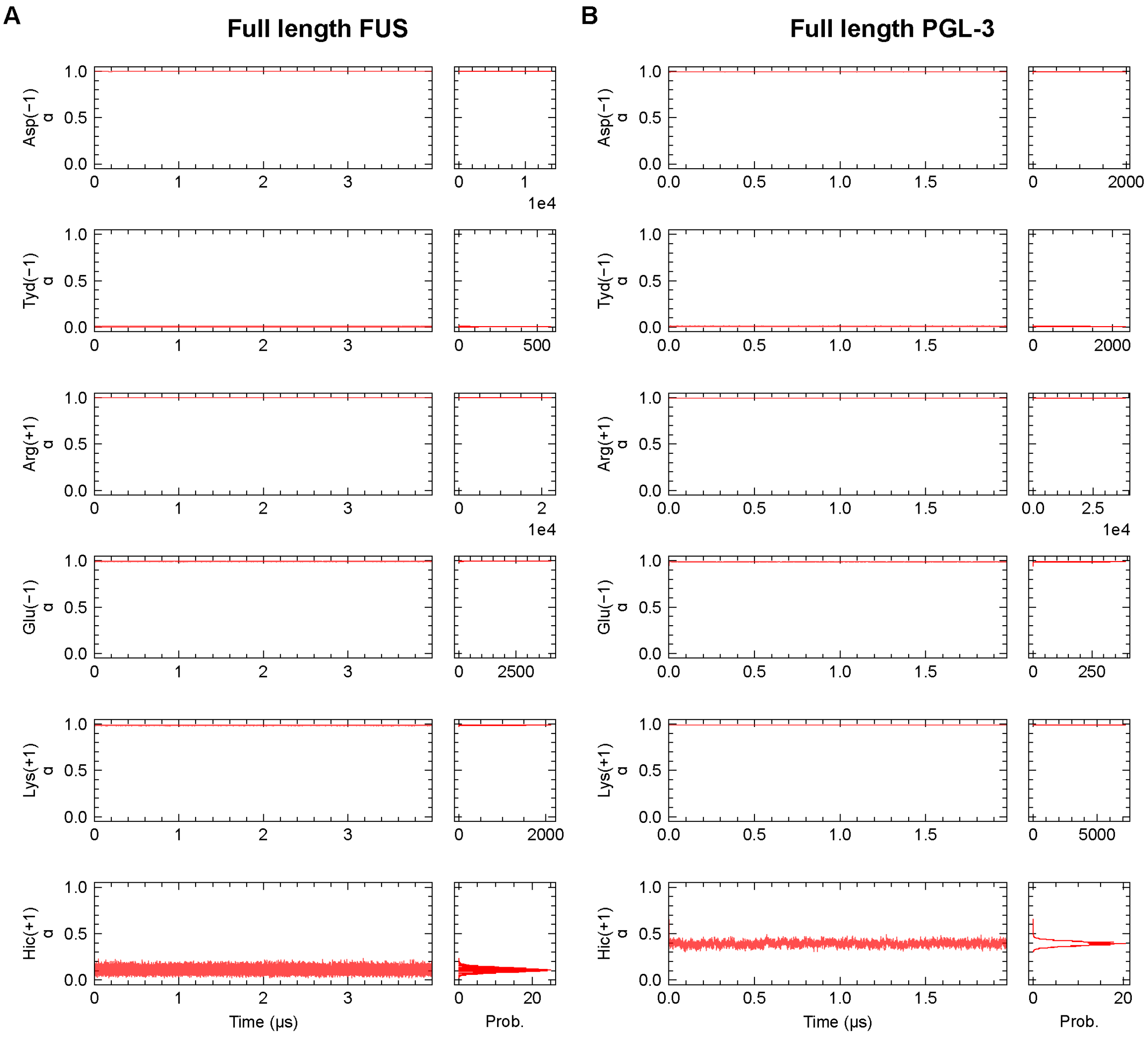


**Fig. S4. Residue-resolved ionization fractions in the dense phase.** Simulation trajectories (left subpanels) and corresponding probability density distributions (right subpanels) of the ionization fractions (α) for all titratable residue types (Asp, Tyr, Arg, Glu, Lys, His), shown for the dense-phase ensembles of **(A)** full-length FUS and **(B)** full-length PGL-3.

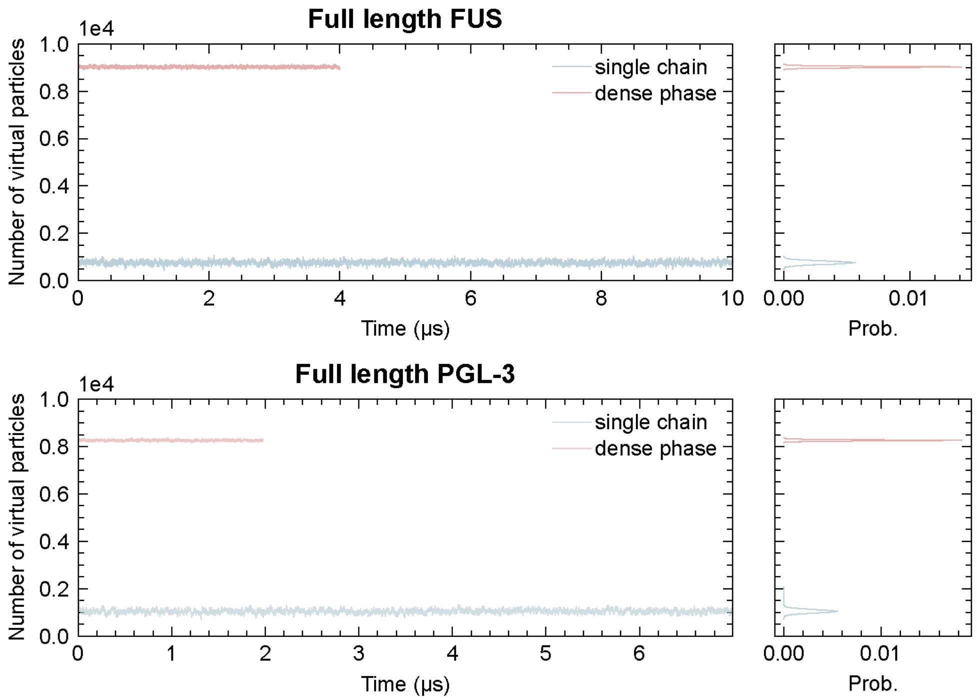


**Fig. S5. Virtual-particle pool occupancy during simulation.** Time evolution (left subpanels) and corresponding probability density distributions (right subpanels) of the virtual-particle count for FL-FUS (top) and FL-PGL-3 (bottom). In all cases, the pool occupancy remained well above zero throughout the entire trajectory, confirming that neither the dilute-phase (single-chain, light blue) nor the dense-phase (pink) simulation approached pool depletion.

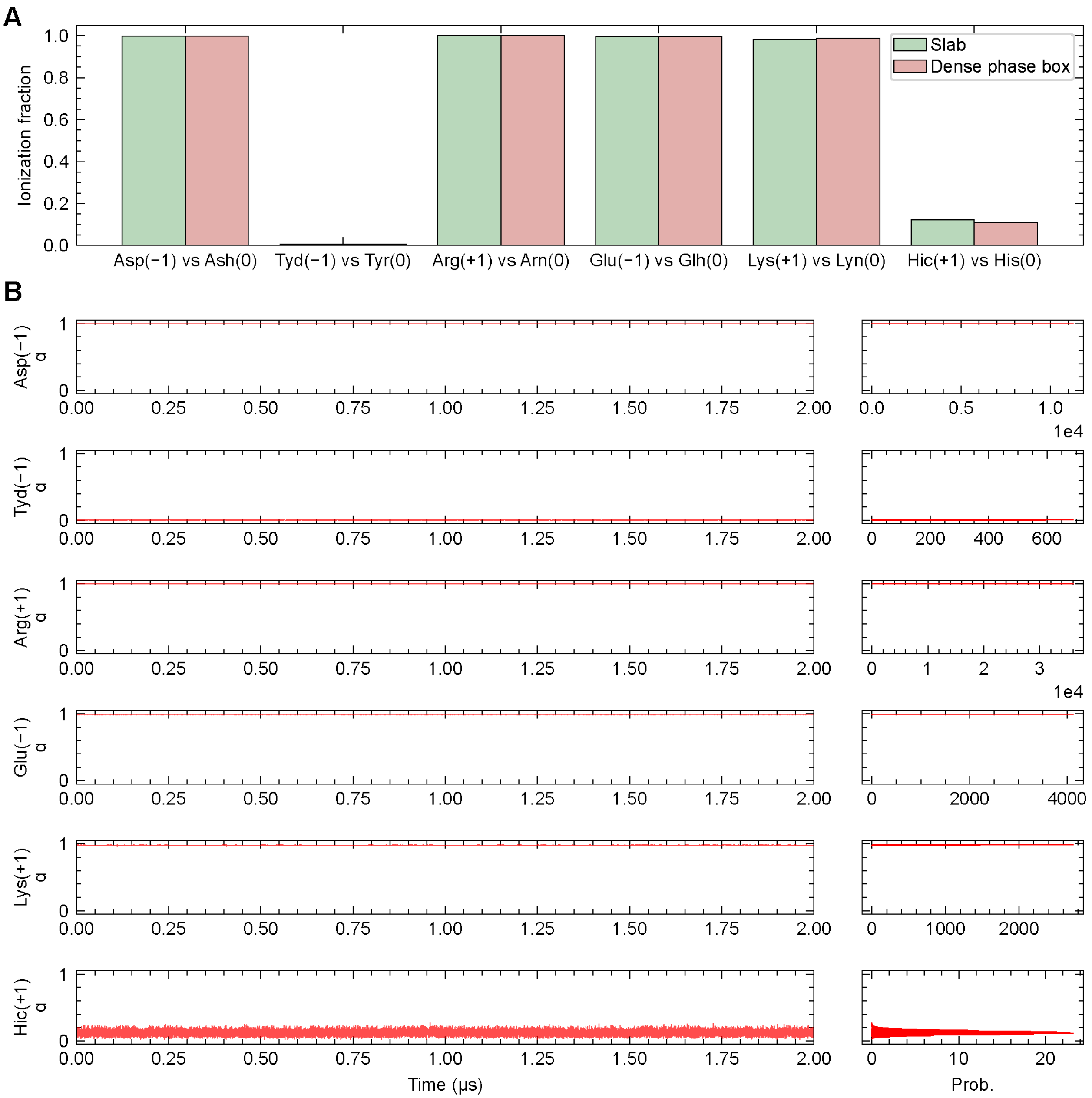


**Fig. S6. Comparison of dense-phase ionization fractions between the slab-geometry and dual-box simulations of FL-FUS. (A)** Mean ionization fractions (α) for all titratable residue types, comparing the slab-geometry simulation (green) with the corresponding dense-phase ensemble of the dual-box protocol (pink). **(B)** Time-resolved trajectories and convergence diagnostics: ionization-state trajectories (left panels) and probability density distributions (right panels), confirming stable sampling and statistically well-converged ionization fractions in both protocols.
